# Three-dimensional chromatin interactions remain stable upon CAG/CTG repeat expansion

**DOI:** 10.1101/754838

**Authors:** Gustavo A. Ruiz Buendía, Marion Leleu, Flavia Marzetta, Ludovica Vanzan, Jennifer Y. Tan, Ana C. Marques, Tuncay Baubec, Rabih Murr, Ioannis Xenarios, Vincent Dion

**Affiliations:** Center for Integrative Genomics, University of Lausanne, 1015 Lausanne, Switzerland; School of Life Sciences, Ecole Polytechnique Fédérale, Lausanne, 1015 Lausanne, Switzerland; Vital-IT Group, Swiss Institute of Bioinformatics, Lausanne, Switzerland; Department of Genetic Medicine and Development, University of Geneva Medical School, Geneva, Switzerland; Institute for Genetics and Genomics in Geneva (iGE3), University of Geneva, Geneva, Switzerland; Department of Computational Biology, University of Lausanne, Lausanne, Switzerland; Department of Molecular Mechanisms of Disease, University of Zurich, Zurich, Switzerland; UK Dementia Research Institute at Cardiff University at Cardiff University, Hadyn Ellis Building, Maindy Road, CF24 4HQ, Cardiff, UK

## Abstract

Expanded CAG/CTG repeats underlie thirteen neurological disorders, including myotonic dystrophy (DM1) and Huntington’s disease (HD). Upon expansion, CAG/CTG repeat loci acquire heterochromatic characteristics. This observation raises the hypothesis that repeat expansion provokes changes to higher order chromatin folding and thereby affects both gene expression in *cis* and the genetic instability of the repeat tract. Here we tested this hypothesis directly by performing 4C sequencing at the *DMPK* and *HTT* loci from DM1 and HD patient-derived cells. Surprisingly, chromatin contacts remain unchanged upon repeat expansion at both loci. This was true for loci with different DNA methylation levels and CTCF binding. Repeat sizes ranging from 15 to 1,700 displayed strikingly similar chromatin interaction profiles. Our findings argue that extensive changes in heterochromatic properties are not enough to alter chromatin folding at expanded CAG/CTG repeat loci. Moreover, the ectopic insertion of an expanded repeat tract did not change three-dimensional chromatin contacts. We conclude that expanded CAG/CTG repeats have little to no effect on chromatin conformation.

## Introduction

The genome is organized into hierarchical topologically associated domains (TADs) (*1*). This three-dimensional organization of chromatin in the nucleus has a profound impact on transcription, DNA replication, recombination, and repair (*2, 3*). For instance, heterochromatic and euchromatic loci are spatially separated and display distinct three-dimensional (3D) chromatin interactions as gleaned by microscopy and chromosome conformation capture (3C)-based techniques (*2, 4, 5*). How this higher-order chromatin structure impinges on biological functions, what determines chromatin domain boundaries, and how it contributes to disease is unclear. Expanded CAG/CTG repeats loci (hereafter referred to according to their mRNA sequence) are ideal to address these questions because they cause diseases and they are associated with changes in local chromatin structure, transcriptional output, and genetic instability (*6*).

CAG/CTG repeats underlie 13 different neurological and neuromuscular disorders including myotonic dystrophy type 1 (DM1) and Huntington’s disease (HD). They are part of a larger group of diseases caused by the expansion of short tandem repeats (STRs) (*7, 8*). Disease-associated STRs (daSTRs) are genetically unstable, especially once they surpass a critical threshold of about 35 to 50 units. Their expansion is associated with extensive chromatin remodeling of the expanded loci (*6, 9-17*). Two such examples are fragile X syndrome (FXS), caused by the expansion of a CGG repeat at the *FMR1* gene located on the X chromosome (*18-21*), and Friedreich’s ataxia (FRDA), caused by a homozygous GAA expansion in the first intron of the *FXN* gene (*22*). In the case of FXS, expansions beyond 200 CGGs are associated with promoter silencing in *cis*. This locus accumulates high levels of heterochromatic marks including CpG methylation, H4K20me3, H3K9me2/3, and H3K27me3 while losing euchromatin-associated marks, such as H3 and H4 acetylation as well as H3K4me2 (*14, 23-26*). In FRDA patient-derived cells, the *FXN* locus with a tendency towards heterochromatinization and loses CTCF binding in cis (*12, 27-30*).

The shift from a euchromatic to a heterochromatic state upon repeat expansion has led to the hypothesis that there is a concurrent change in higher order chromatin folding. Indeed, 3C-based experiments have revealed an increase in the frequencies of 3D chromatin interactions surrounding expanded GAA and CGG repeats in patient-derived lymphoblastoid cell lines (LCLs) (*12, 23, 31*). This was interpreted as enhanced compaction around the *FXN* locus and as a rearrangement of chromatin contact domains at the *FMR1* locus in FXS cells, respectively. It was speculated that higher order chromatin folding might contribute to the silencing of the genes in the vicinity of the expanded repeat (*12, 23, 31*). Importantly, daSTRs are found predominantly at TAD and sub-TAD boundaries, suggesting more generally that daSTR expansions may disrupt TADs (*31*). This may then contribute to gene silencing in cis and to the high levels of instability found at these sequences, ultimately altering disease progression (*31*).

The critical unknowns in this model are whether changes in higher-order chromatin structure are confined to CGG and GAA repeats or if this is general to daSTRs and whether changes in chromatin interactions cause alterations in gene expression and repeat instability. The later hypothesis is especially appealing in the context of the expanded CTG repeats in the 3’UTR of the *DMPK* gene in DM1 cells because, like expanded CGG and GAA repeat loci, this region undergoes heterochromatinization upon repeat expansion. The changes observed at the *DMPK* locus include the loss of CTCF binding and of a DNAseI hypersensitive site, an increase in DNA and H3K9 methylation, as well as a loss of acetylated histone marks around the repeat tract (*13, 16, 17, 32-34*). Moreover, the ectopic introduction of an expanded CAG repeat in yeast, was sufficient to relocate the locus to the nuclear periphery in S-phase budding yeast cells leading to changes in repeat size (*35*). Importantly, expanded CAG/CTG repeats cause the majority of daSTR disorders. Together, these observations prompted us to test the hypothesis that chromatin conformation changes at expanded CAG/CTG repeats. We used 4C-seq in LCLs from DM1 and HD individuals and in HEK293-derived cells harboring an ectopic CAG repeat expansion. We found no evidence to support changes in chromatin interactions that would underlie the alterations in transcriptional output or genetic instability of these sequences. This was consistent in different cell types, genetic backgrounds, and in the presence of specific heterochromatin marks. We conclude that changes in higher order chromatin folding do not contribute to the pathogenesis of expanded CAG/CTG repeat disorders.

## Results

### Chromatin conformation is stable upon repeat expansion at the *HTT* locus

To assess whether chromatin contacts change upon CAG repeat expansion at the *HTT* locus, we used a series of HD patient-derived LCLs (GM02164, GM03620, and GM14044, referred to as HD-A, HD-B, HD-C respectively) as well as two lines from unaffected individuals (GM04604 and GM02180, UN-A and UN-B respectively). Their family relationships and their repeat sizes are found in Fig. 1A, S1A, and Table S1.

**Fig. 1.**
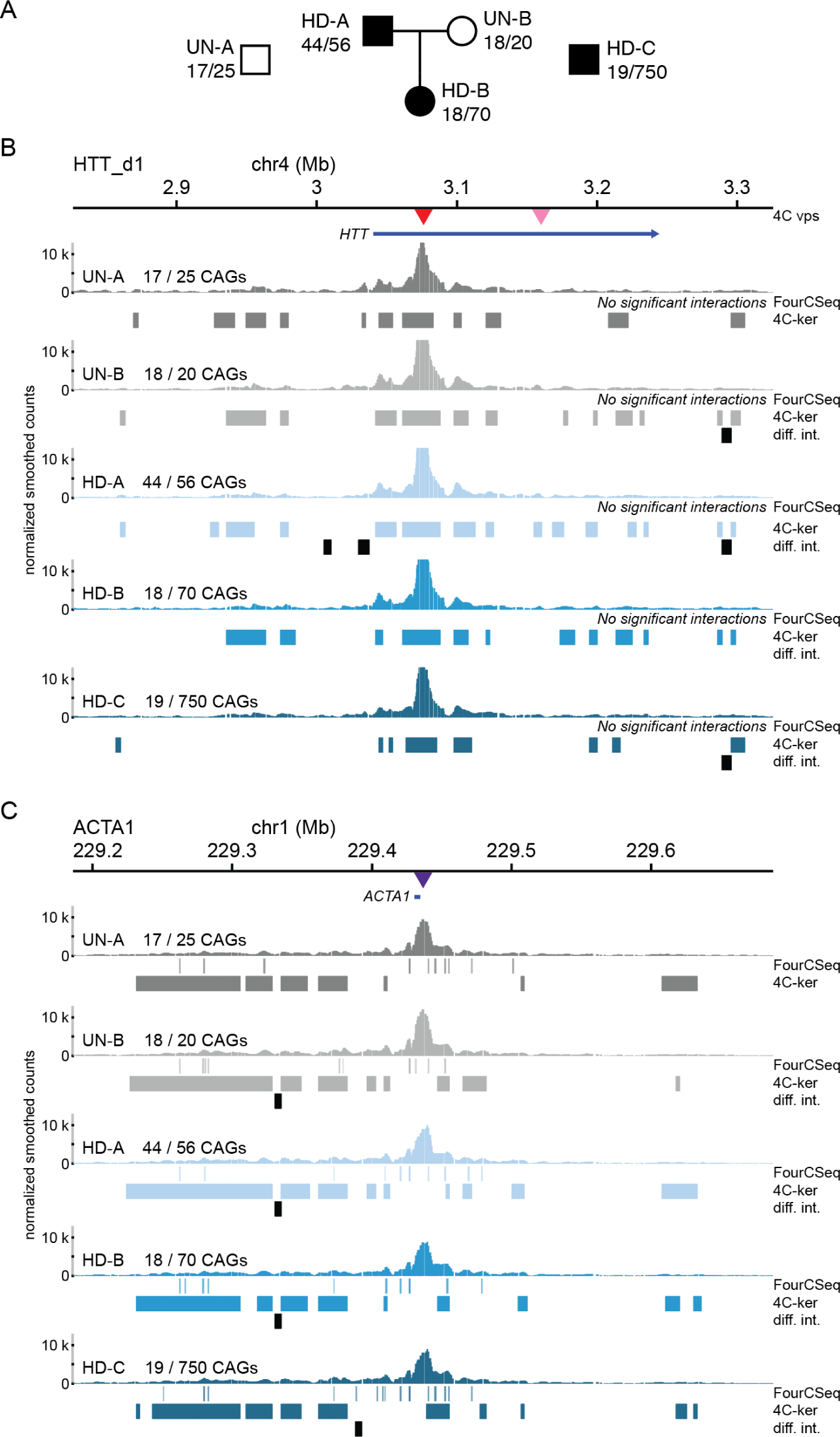
Chromatin interactions of the *HTT* locus in unaffected and HD cells. (A) Pedigree of HD patient cell lines used in this study along with unrelated unaffected and HD cell lines (UN-A and HD-C respectively). **(B)** 4C-seq chromatin interaction profiles (average of triplicate smoothed and normalized counts) from the HTT_d1 viewpoint (1 kb downstream of the CAG repeats - red central triangle) in two unaffected (UN-A and UN-B) and three HD patient cell lines (HD_A, HD_B, and HD_C). The top blue bar represents the *HTT* gene and the triangles represent the location of both *HTT* viewpoints. The interaction profiles for the HTT_d85 viewpoint (85 kb downstream of the CAG repeat) can be found in Fig. S3. **(C)** 4C-seq chromatin interaction profiles (average of triplicate smoothed and normalized counts) from the *ACTA1* viewpoint (central purple triangle). For panels B and C, high-interacting regions were called using 4C-ker and significant interactions were called using FourCSeq. Regions of differential interactions are marked with black bars below each 4C-seq track and labeled as “diff. int.”. The top blue bar represents the *ACTA1* gene.

To determine chromatin conformation, we used 4C-seq (*36, 37*) because it maximizes resolution at the loci of interest. This allows a high sensitivity to small changes in conformation that may be missed by other 3C-based methods. For the *HTT* locus, we used 2 viewpoints on chromosome 4 within the gene body – 1 kb (HTT_d1) and 85 kb (HTT_d85) downstream of the CAG repeats. To control for a potential effect of the pathology on genome-wide chromatin conformation, we also used a viewpoint located near the *ACTA1* gene on chromosome 1. We obtained three replicates of each 4C viewpoint and compared the DNA interaction profiles between the unaffected and HD cell lines (Fig. 1B). Replicates from the same cell lines show good correlation in fragments with more than 20 mapped reads (Fig. S2A-C).

We then identified fragments and regions that interact with the 4C viewpoint at frequencies higher than expected (significant interactions, see methods). We also looked for regions that show significant differences in interaction frequencies between patient-derived and unaffected cells (differential interactions, see methods). To do so, we used two 4C-seq data analysis packages: FourCSeq (*38*) and 4C-ker (*39*). We found that the interaction profiles were similar within a 2 Mb region around the viewpoint (Fig. 1B, Fig. S3). Notably, the chromatin conformation remained unaltered in HD-A, an HD patient cell line that has two expanded alleles (44/56 CAGs), as well as in HD-B (18/70 CAGs) and HD-C (19/750 CAGs). The *ACTA1* viewpoint also produced indistinguishable interaction profiles between unaffected and HD patient LCLs (Fig. 1C). We identified few small regions displaying differential interactions, but they were mainly outside of regions with significant interactions (Fig. 1BC). In addition, most regions of differential interactions were not exclusive to HD patient cells, as they were also found in comparisons between the two unaffected cell lines. This suggests that the minor changes in interaction frequencies are due to factors other than the presence of the expanded repeat tract in *HTT*, e.g. the different genetic backgrounds. Taken together, these results show that a CAG repeat expansion at the *HTT* locus does not cause significant alterations in chromatin conformation.

### Chromatin conformation is stable upon repeat expansion at the *DMPK* locus

Repeat expansion at the *HTT* locus is not known to be associated with significant changes in histone marks and chromatin accessibility, which might cause changes in chromatin conformation. To determine whether our findings applied in a case where chromatin structure is altered around expanded CAG/CTG repeats, we studied four viewpoints in the *DMPK* gene. We used two DM1 patient-derived LCLs (GM06077 and GM04648, DM1-A and DM1-B respectively). The DM1-A cell line harbored one expanded *DMPK* allele with 1,700 CTGs and the DM1-B cell line, 1,000 CTGs (Fig. 2A and S1B). We found that DM1-A cells had increased CpG methylation levels at two CTCF binding sites flanking the repeats (Fig. S4B), with a concomitant loss of CTCF binding at these sites (Fig. S4C). In DM1-B cells we observed normal methylation levels at both CTCF binding sites and slightly reduced CTCF binding (Fig S4B). Thus, DM1-A cells displayed molecular signatures of congenital DM1 (*40*), whereas DM1-B cells have adult onset DM1 characteristics (Table S1).

**Fig. 2.**
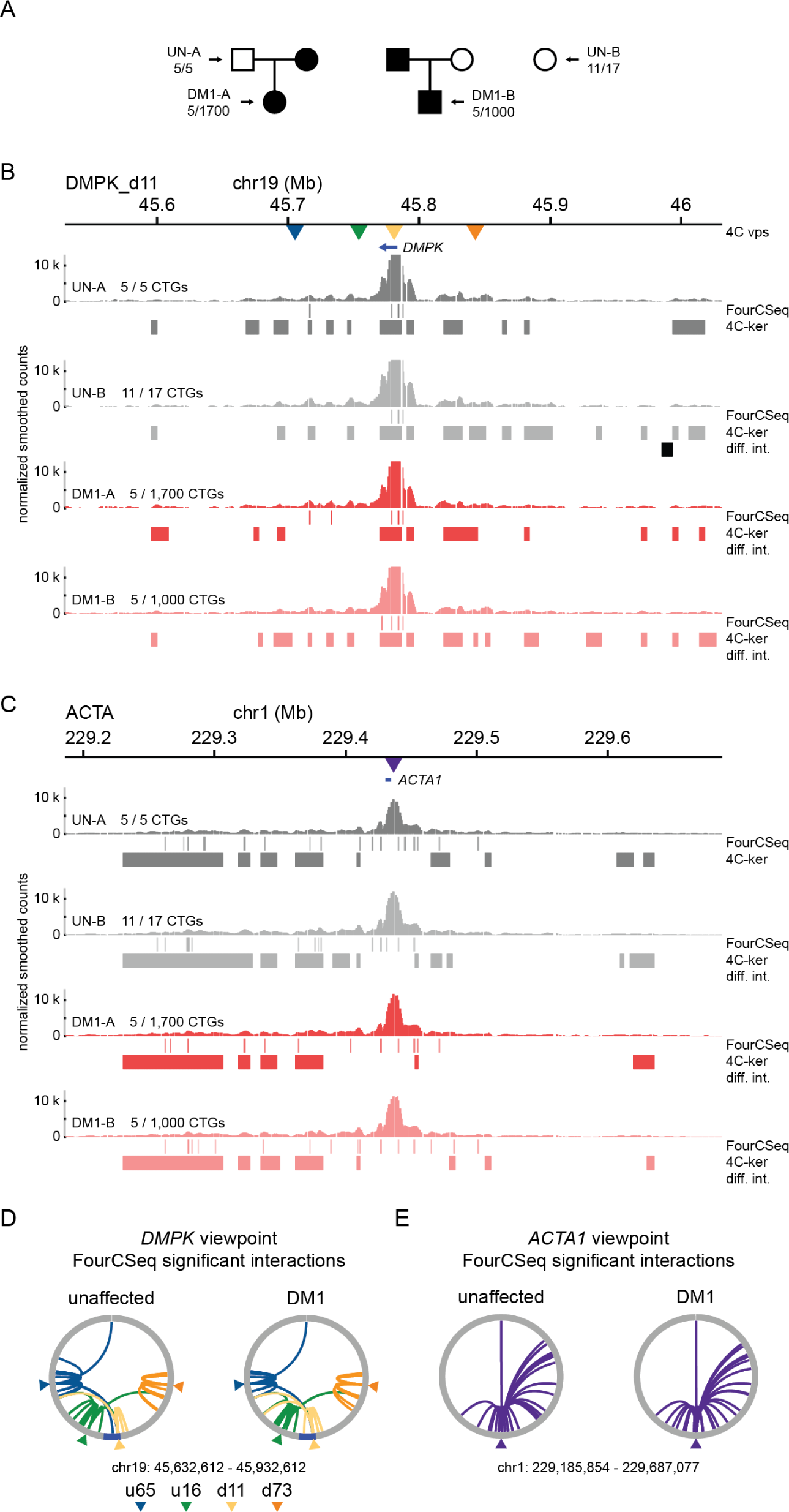
Chromatin interactions of the *DMPK* locus in unaffected and DM1 cells. **(A)** Pedigree of DM1 patient cell lines used in this study along with an unrelated unaffected cell line (UN-B). **(B)** 4C-seq chromatin interaction profiles (average of triplicate smoothed and normalized counts) from the DMPK_d11 viewpoint (11kb downstream of the CTG repeats - yellow triangle) in two unaffected (UN-A and UN-B) and two DM1 patient cell lines (DM1-A and DM1-B). The top blue bar represents the *DMPK* gene and the triangles represent the location of the four *DMPK* viewpoints. The profiles for the three other viewpoints can be found in Fig. S5. **(C)** 4C-seq chromatin interaction profiles (average of triplicate smoothed and normalized counts) from the *ACTA1* viewpoint (central purple triangle) in two unaffected and two DM1 patient cell lines. The top blue bar represents the *ACTA1* gene. For panels B and C, high-interacting regions were called using 4C-ker and significant interactions were called using FourCSeq. Regions of differential interactions are marked with black bars below each 4C-seq track and labeled as “diff. int.”. **(D)** Circos plot of the significant interactions called with FourCSeq (nominal p-value < .05) from four different viewpoints surrounding the CTG repeats of *DMPK* in the unaffected and DM1 cell lines (left and right respectively) (DMPK_u65 in blue, DMPK_u16 in green, DMPK_d11 in yellow, and DMPK_d73 in orange, 65 kb upstream, 16 kb upstream, 11 kb downstream, and 73 kb downstream, respectively). **(E)** Circos plot of the significant interactions called with FourCSeq (nominal p-value < .05) from the control *ACTA1* viewpoint in unaffected and DM1 cell lines (left and right respectively).

We performed 4C-seq on the unaffected and DM1 cell lines using four different 4C viewpoints at distinct distances away from the *DMPK* CTG repeats (Fig. 2A). Replicates from the same cell lines also showed good correlation in 4C fragment counts above 20 (Fig. S2C-G). Similar to the HD scenario, we observed strikingly similar chromatin interaction profiles between unaffected and DM1 samples for all four viewpoints in the *DMPK* region (Fig. 2B, Fig. S5). As expected, the *ACTA1* viewpoint showed interaction profiles indistinguishable between unaffected and DM1 cell lines (Fig. 2C). As with the *HTT* viewpoints, none of the *DMPK* or *ACTA1* viewpoints in DM1 patient cells had significant interactions that were also called as regions of differential interaction (Fig. 2B-E). Taken together, we found no evidence for large-scale changes in chromatin interactions at the *DMPK* locus driven by CTG expansions, despite the changes in CTCF occupancy and DNA methylation levels.

### Lack of allelic bias in chromatin interactions at expanded CAG/CTG repeats

DM1 and HD are both dominantly inherited disorders with individuals being heterozygous for the expanded allele. Thus, one potential caveat in our data was that the presence of a normal-length allele masks changes in 3D chromatin interactions made by the expanded allele. To evaluate this possibility, we took advantage of the presence of at least one parental cell line for the DM1-A and HD-B patient cell lines in our dataset (Fig. 1A and 2A) and identified allele-specific single nucleotide polymorphisms (SNPs) in the 4C-seq data from these samples. We selected a subset of the SNPs that could be assigned unambiguously to either the expanded or the normal allele within 1 Mb of the 4C viewpoints (see methods). We reasoned that if chromatin contacts were established without a systematic bias for either allele in DM1 and HD patient cells, the proportion of 4C fragments in which the expanded chromosome had more reads than the normal-length one would be close to 50%. We analyzed the sequencing coverage of the 4C data at these SNPs positions and found that the viewpoints did not establish preferential contacts with a single chromosome in neither the DM1-A nor HD-B patient cell lines (Table 1). These results are consistent with the conclusion that chromatin interactions at both disease loci do not show allelic bias. Together, these results corroborate the conclusion that expanded CAG/CTG repeats do not significantly alter the DNA interactions at two expanded CAG/CTG repeat loci.

**Table 1.** Allele-specific interactions in 4C-seq data.

| Cell line | Disease | Genomic region <sup>1</sup> | Total # of samples | # of samples with more normal chromosome reads | # of samples with more expanded chromosome reads | p-value <sup>2</sup> |
| --- | --- | --- | --- | --- | --- | --- |
| DM1-A | DM1 | *ACTA1* | 6 | 1 | 5 | 0.22 |
|  |  | *DMPK* | 28 * | 18 | 9 | 0.09 |
| HD-B | HD | *ACTA1* | 5 | 2 | 3 | 1 |
|  |  | *HTT* | 19 | 7 | 12 | 0.25 |

<sup>1</sup> 1Mb region around the 3'end of *DMPK*, 5' end of *HTT*, or the *ACTA1* 4C-seq viewpoint.
<sup>2</sup> p-value of an exact binomial test.
\* 1 sample had the same number of reads in both alleles, thus it was not included

### Chromatin conformation at an ectopic CAG repeat locus

It remained possible that the genetic background may have had a confounding effect on the chromatin interactions made at expanded CAG/CTG repeat loci. To test this, we compared the chromatin interactions of a hemizygous ectopic locus with either normal-length or expanded CAG/CTG repeats in otherwise isogenic cell lines. We obtained two clonal populations of Flp-In T-REx 293 cells that contain a single, stably integrated construct containing CAG repeats within the intron of a GFP mini-gene (*41, 42*). One cell line contained 15 CAGs (GFP(CAG)15) whereas the other had 270 CAGs (GFP(CAG)270) which is within the pathogenic range for both DM1 and HD. The reporter sports a doxycycline-inducible promoter. Using targeted locus amplification (*43*) and TAIL PCR, we mapped the insertion site to the p-arm of chromosome 12, 1.2 Mb from the telomere (Fig 3A). We performed 4C-seq in both cell lines with a viewpoint located 1 kb upstream of the CAG repeats. The chromatin interaction profiles of this ectopic CAG locus were also very similar between the 15 and 270 CAG repeat cells, with few regions of differential interactions overlapping with those displaying significant interactions using 4Cker. By contrast, FourCSeq did not detect any changes that were significant (Fig. 3B). We concluded that CAG repeats at an ectopic site causes few changes to chromatin conformation.

**Fig. 3.**
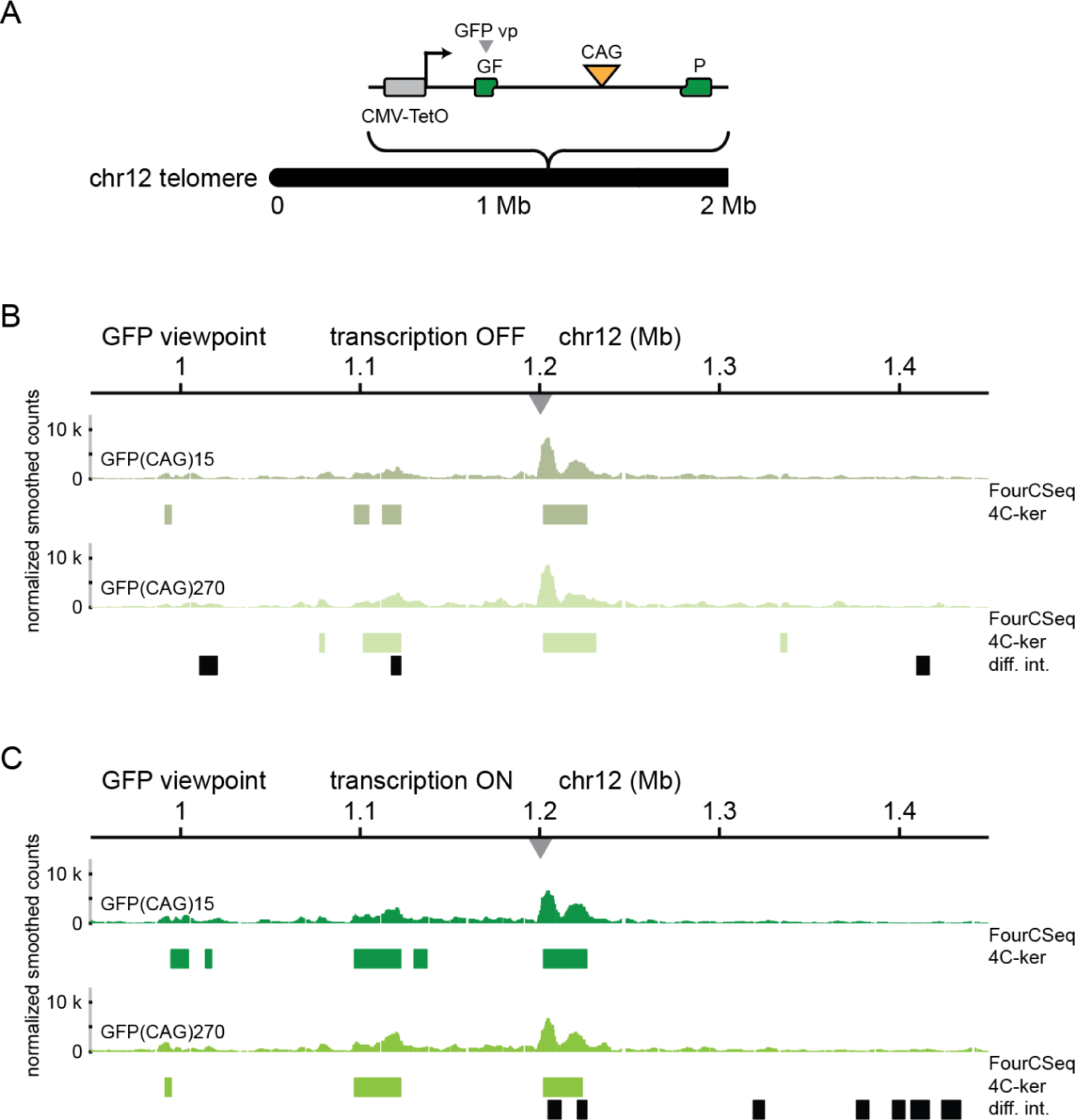
Chromatin interactions of an expanded CAG repeat locus in isogenic cells. **(A)** Diagram of the integration site of the CAG ectopic locus in GFP(CAG)n cells. **(B)** 4C-seq chromatin interaction profiles (average of triplicate smoothed and normalized counts) from the GFP viewpoint in GFP(CAG)15 (top) and GFP(CAG)270 (bottom). Cells were treated with doxycycline to induce transcription of the GFP mini-gene. **(C)** 4C-seq chromatin interactions from the GFP viewpoint in GFP(CAG)15 (top) and GFP(CAG)270 (top) when transcription was not induced.

Some studies suggest that transcription may help define topological domain boundaries (*44-47*). To determine whether transcription through expanded CAG repeats could lead to changes in chromosome conformation, we induced transcription by culturing the GFP(CAG)n cells with doxycycline for five days. We observed that only 2 of 17 regions of differential interactions overlapped fully or in part with high-interacting regions over a 2 Mb region using 4Cker. FourCSeq, on the other hand, found no significant interactions (Fig. 3C). These results suggest that an ectopic expanded CAG repeat tract is not enough to alter the chromosome conformation significantly at this locus, regardless of the transcriptional status.

## Discussion

Here we showed that chromatin interactions remain stable at expanded CAG/CTG repeat loci. This was true for two disease loci and one transgene and in two different cell types. Our findings are also supported by allele-specific analysis of 4C-seq interactions. Furthermore, CpG methylation and CTCF binding at two sites flanking the CTG repeats of *DMPK* did not impact chromosome conformation. This is especially relevant because CTCF is a key architectural protein involved in the demarcation of TAD and sub-TAD boundaries (*48*). Additionally, when we inserted a hemizygous transgene with CAG repeats, we found that expanded CAG repeats did not induce a significant re-organization of the chromatin contacts established at this exogenous CAG/CTG repeat locus. These results further show that an expanded CAG/CTG repeat tract is not sufficient to change chromatin folding.

One possibility is that expanded CAG/CTG repeats lead to changes in chromatin conformation in specific cell types that are especially vulnerable in DM1 or HD, for example cardiomyocytes or medium spiny neurons, but not in LCLs. However, that would complicate the hypothesis by adding a requirement for a cell-type specific factors to mediate these changes. If such a factor exists, our conclusion remains unaffectcted: that the expansion of CAG/CTG repeats is not enough to effect changes in chromatin conformation.

Our findings are in contrast to the effect of expanded CGG and GAA repeats on chromosome conformation (*12, 23, 31*). In FXS patient-derived B-lymphoblast and fibroblasts, expanded CGG repeats in *FMR1* were correlated with heterochromatic characteristics, decreased CTCF binding, and ultimately a disruption of a TAD boundary near the expanded repeats (*31*). The DM1 patient cells used here, especially the congenital DM1 patient cell line, share similar changes in chromatin marks to FXS patient cells. And yet, these factors did not amount to an alteration of the 3D chromatin interactions at the expanded *DMPK* locus. There are important differences between FXS/FRDA and DM1/HD that could account for the differences. Apart from the nucleotide composition of the repeat tract themselves, the disease loci and thus their flanking sequences are different. Further investigations are needed to understand what accounts for the differences.

It was speculated that the changes in chromatin conformation found at expanded CGG repeats could lead to changes in repeat instability (*31*). Indeed, CGG repeat expansions beyond 200 units are associated with promoter silencing, chromatin contact alterations and their instability is lessened (*49, 50*). By contrast, long GAA and CAG/CTG repeats are more unstable (*7, 51, 52*), yet the chromosome conformation at expanded GAA repeats is altered but not at expanded CAG/CTG repeats. Thus, there does not seem to be a correlation between changes in 3D chromatin contacts and repeat instability. One possibility is that the effect of chromosome conformation on repeat instability is repeat-type specific. Another is that chromosome conformation is unrelated to the repeat instability and is neither a cause nor a consequence. Similarly, changes in transcriptional output around expanded repeats may be unrelated to chromatin conformation and rather be the consequence of changes in local chromatin marks, or they may be specific to the repeat sequence.

The expanded CAG/CTG repeat loci in DM1 and HD provide an endogenous genomic substrate to study the mechanisms necessary for the establishment of 3D chromatin domains in daSTRs given the involvement of CTCF binding, CpG methylation, and other chromatin remodeling events at these loci. Although CTCF is generally a major driver of genome organization, it is not the only driver. There are cases where CTCF binding abrogation does not alter an evolutionarily conserved TAD boundary, for instance at the *Firre* locus (*53*). Moreover, our results are in line with recent evidence in *Drosophila* arguing that genome topology is not predictive of transcriptional output genome-wide (*54*). Indeed, our data argue that changes in chromatin topology are unlikely to underpin the molecular pathology of expanded CAG/CTG repeat disorders.

## Materials and Methods

### Cell lines

All LCLs were obtained from the Coriell Institute for Medical Research Cell Repository. They were grown at 37°C and 5% CO2 in RPMI 1640 medium supplemented with 15% FBS, 2 mM L-glutamine, and 1% penicillin-streptomycin. Cells were counted and passaged every 3-4 days depending on cell density. The GFP(CAG)270 and GFP(CAG)15 lines were previously characterized (*41*). They were maintained at 37°C and 5% CO2 in DMEM with Glutamax, 1% penicillin-streptomycin, 15 μg mL^−1^ blasticidine, and 150 μg mL^−1^ hygromycin. Transcription of the GFP mini-gene was activated by culturing GFP(CAG)n cells with doxycycline at a final concentration of 2 µg mL^-1^ for five days.

### Repeat length determination and small-pool PCR

Genomic DNA was isolated from each LCL using the NucleoSpin Tissue kit (Macherey-Nagel). PCR products containing the CTG repeats from *DMPK* were amplified with primers oVIN-1252 and oVIN-1251 (Table S3). PCR products with the CAG repeats from *HTT* were produced with primers oVIN-1333 and oVIN-1334 (Table S3). For normal-length alleles, several PCRs were set up with Mango Taq (Bioline) and the products were gel-extracted and Sanger-sequenced with the same primers used for the amplification. For expanded alleles, small-pool PCR was performed based on a previously described protocol (*55*). Briefly, the same primers were used for the amplification of expanded *DMPK* and *HTT* alleles with 1 ng of genomic DNA per PCR. The products were run on an agarose gel and transferred to a nylon membrane. An oligo made up of 10 CAGs was used to obtain a radioactive probe used for the visualization of the expanded alleles. The number of repeats reported here is an estimation of the modal number of repeats.

### Bisulfite sequencing

Bisulfite sequencing was performed according to a previously described method (*40*). Bisulfite conversion was performed with the EZ DNA Methylation kit (Zymo Research, D5001) using the standard protocol. 200 ng of genomic DNA were used for bisulfite conversion at 50°C for 12 hours.

Bisulfite-converted DNA was desulphonated, eluted, and immediately used for nested and hemi-nested PCR amplification of the upstream and downstream CTCF binding sites respectively using previously described primers (*40*). 50 ng of bisulfite-converted DNA were used for the first PCR, and 3 µL of the products were used for the second PCR. The final amplicons were purified with the NucleoSpin PCR Clean-up kit (Macherey-Nagel, 740609) and used for 2×250 bp paired-end sequencing (Illumina). The primers used for both rounds of PCR are found in Table S3. Sequencing reads were preprocessed using TrimGalore (github.com/FelixKrueger/TrimGalore) with the following parameters: -q 20 --length 20 –paired. Reads were aligned using QuasR (Gaidatzis et al 2015, PMID 25417205) to human reference (GRCh38) DNA sequences corresponding to the amplified PCR products. DNA methylation calls for each CpG were extracted using the qMeth() function in QuasR. DNA methylation frequencies were calculated as methylated CpGs / total covered CpG x 100 and plotted in R.

### CTCF ChIP-qPCR

Chromatin immunoprecipitation was performed according to the Diagenode AutoiDeal ChIP-qPCR kit (Diagenode, C01010181) standard protocol. Samples of 6×10^6^ cells were sonicated using the Diagenode Bioruptor Pico (Diagenode, B01060010), with 10 cycles of 30 sec “on” and 30 sec “off”. Correct DNA fragmentation was verified by agarose gel electrophoresis. Immunoprecipitation was performed with 4×10^6^ cells using a CTCF antibody (Diagenode, C15410210) and the Diagenode IP Star Compact Automated System robot (Diagenode B03000002). Results were analyzed using the StepOnePlus qPCR by Applied Biosystems (ThermoFisher Scientific, 4376357) with Applied Biosystems SYBR^TM^ Green PCR Mastermix (ThermoFisher Scientific, 4309155). Primer sequences for qPCR are listed in Table S3.

### 4C-seq

4C library preparation was performed based on a previously described protocol (*56*). 10^7^ cells were crosslinked in 2% formaldehyde for 10 minutes at room temperature and quenched with glycine to a final concentration 0.13 M. Cells were lysed for 15 minutes on ice in lysis buffer (50 mM Tris-HCl pH 7.5, 150 mM NaCl, 5 mM EDTA, 0.5% NP-40, 1% Triton X-100). The first digestion was performed at 37°C overnight with 600 U of DpnII (NEB) followed by DpnII heat inactivation. The first ligation was performed at 16°C overnight with 50 U T4 DNA ligase (Thermo Scientific) in 7 mL. Ligated samples were de-crosslinked with 30 µL Proteinase K (10 mg/mL) at 65°C overnight followed RNAse A treatment and phenol-chloroform purification. The second digestion was performed with 50 U of BfaI (NEB) at 37°C overnight followed by BfaI heat inactivation. The second ligation was performed at 16°C overnight with 100 U T4 DNA ligase in 14 mL. 4C template samples were purified with phenol-chloroform and QIAquick PCR purification kit columns. 4C libraries were generated by amplifying 1 µg total of purified 4C template with 4C viewpoint primers, pooling reactions and purifying the PCR products with AMPure XP beads to exclude products less than 130 bp. Single-end sequencing of pooled 4C libraries was performed on Illumina HiSeq 2500.

### 4C-seq data analysis

Demultiplexing and mapping were performed using the BBCF HTSstation (*57*) according to (*58*). The 4C fragments surrounding the viewpoints (±2.5 kb) were excluded from the rest of the analysis. The demultiplexed reads were mapped to the human genome GRCh38 using bowtie2 (v 2.2) (*59*). Fragment counts were obtained using FourCSeq (v1.18.0) (*38*). The number of mapped reads for each sample is found in Table S2. For plotting the data, fragment counts were normalized (reads per million) and smoothed with a running mean (window size = 5 fragments). The smoothed and normalized fragment counts were averaged among replicates of the same 4C library samples and visualized with gFeatBrowser (http://www.gfeatbrowser.com). For the FourCSeq analysis, we defined significant interactions as fragments with a z-score equal to or greater than 1.96 and a false-discovery rate of 0.1, using the following parameters: minCount=20 and fitFun=“distFitMonotone” in getZScores function; zScoreThresh=1.96, fdrThresh = 0.1 in addPeaks function. 4C-ker (v0.0.0.9000) uses a Hidder-Markov Model that accounts for differences in coverage near 4C viewpoints to determine three types of domains: high-interacting, low-interacting, and non-interacting domains. For each viewpoint, we used k=5 in the nearBaitAnalysis function, and plotted the high-interacting regions. The difference between significant interactions called with FourCSeq and 4C-ker is expected given that FourCSeq usually identifies “peaks” of significantly interacting regions whereas 4C-ker identifies regions (*60*). The analysis of differential interactions was performed with the differentialAnalysis function of 4C-ker, which is based on the DESeq2 framework, using default parameters (including a p-value threshold of 0.5). The UN-A cell line was used as the reference condition for all comparisons.

### 4C SNP data analysis

We called SNPs from 4C-seq data from the DM1-A and HD-B patient-derived samples using GATK (v3.7.0) (*61*) and samtools/bcftools (v1.5 and v1.4.1 respectively) (*62*). Biallelic SNPs located within a 1 Mb region of the 3’ end of *DMPK*, the 5’ end of *HTT*, and the *ACTA1* 4C viewpoint were selected. Among these, only SNPs that could unambiguously inform which parental allele they came from were retained for downstream analysis. This required a homozygous genotype in at least one of the parents’ cell lines. We validated a subset of these SNPs by PCR amplification of the genomic region encompassing the variants followed by library preparation and paired-end sequencing with Illumina Miseq nano 2 x 250 bp. We then analyzed the sequencing coverage at allele-specific SNP positions from the 4C-seq data from DM1-A and HD-B patient cell lines. For samples with at least 10 reads per SNP position, we counted the number of times the expanded allele had more mapped reads than the normal allele. We applied an exact binomial test to statistically assess whether the proportion of cases where the expanded allele had more reads than the normal allele was significantly different to 0.5, which represented the null hypothesis of no allelic bias.

### Targeted Locus Amplification

Targeted locus amplification was performed based on a previously described protocol (*43*). 10^7^ cells were crosslinked in 2% formaldehyde for 10 minutes at room temperature and quenched with glycine to a final concentration of 275 mM. Cell were lysed for 5 minutes at room temperature in lysis buffer (50 mM Tris-HCl pH 7.5, 150 mM NaCl, 5 mM EDTA, 0.5% NP-40, 1% Triton X-100). Crosslinked samples were digested at 37°C overnight with 400 U NlaIII (NEB) followed by NlaIII inactivation. Samples were ligated at room temperature for 2 hours with 20 U T4 DNA ligase (Thermo Scientific) in 500 µL. Ligated samples were de-crosslinked with 5 µL Proteinase K (10 mg/mL) at 65°C overnight followed by RNAse A treatment and phenol-chloroform purification. Samples then digested overnight at 37°C with 50 U NspI (NEB) followed by NspI inactivation. The second ligation was performed overnight at 16°C with 100 U T4 DNA ligase in 14 mL. TLA circularized templates were purified with QIAquick PCR purification kit columns. TLA libraries were generated by amplifying 800 ng of purified TLA template with TLA viewpoint primers. Paired-end sequencing of pooled TLA libraries was performed on Illumina HiSeq 4000. Reads were mapped using a custom TLA analysis pipeline using the BWA mapping software (v0.7.17) (*63*). First, reads were mapped to the human genome GRCh38. Then, unaligned sequences are digested *in-silico* with the NlaIII restriction site and remapped to the genome. The combined mapping results were used to determine the integration site.

## Acknowledgments

We thank Emma Randall for the sequencing of the *HTT* repeats for the UN-A and UN-B cells as well as the *DMPK* repeats of UN-B cells. We thank Fisun Hamaratoglu for critical reading of the manuscript, Frédéric Schütz for statistical counsel, and Andrzej Stasiak for scientific discussions. The high throughput sequencing was performed by the Genomics Technology Facility at the University of Lausanne.

## Author contributions

G.A.R.B. and V.D. designed the experiments, prepared the figures, and wrote the manuscript with input from all the authors. G.A.R.B. performed the experiments and data analysis. M.L. and G.A.R.B. performed the analysis of the 4C-Seq data. F.M. performed the SNP calling and G.A.R.B. performed the statistical analysis. J.Y.T. and A.C.M. helped with mapping the insertion site. L.V. and R.M. performed the CTCF ChIP-qPCR. G.A.R.B. performed the bisulfite sequencing experiments and T.B. analyzed the data. I.X. supervised G.A.R.B. together with V.D. and helped in designing the experiments.

## Data and material availability

All sequencing data underlying this study will be shortly deposited in the SRA of NCBI. All data presented here are available from the corresponding author.

---

**Fig. S1.**
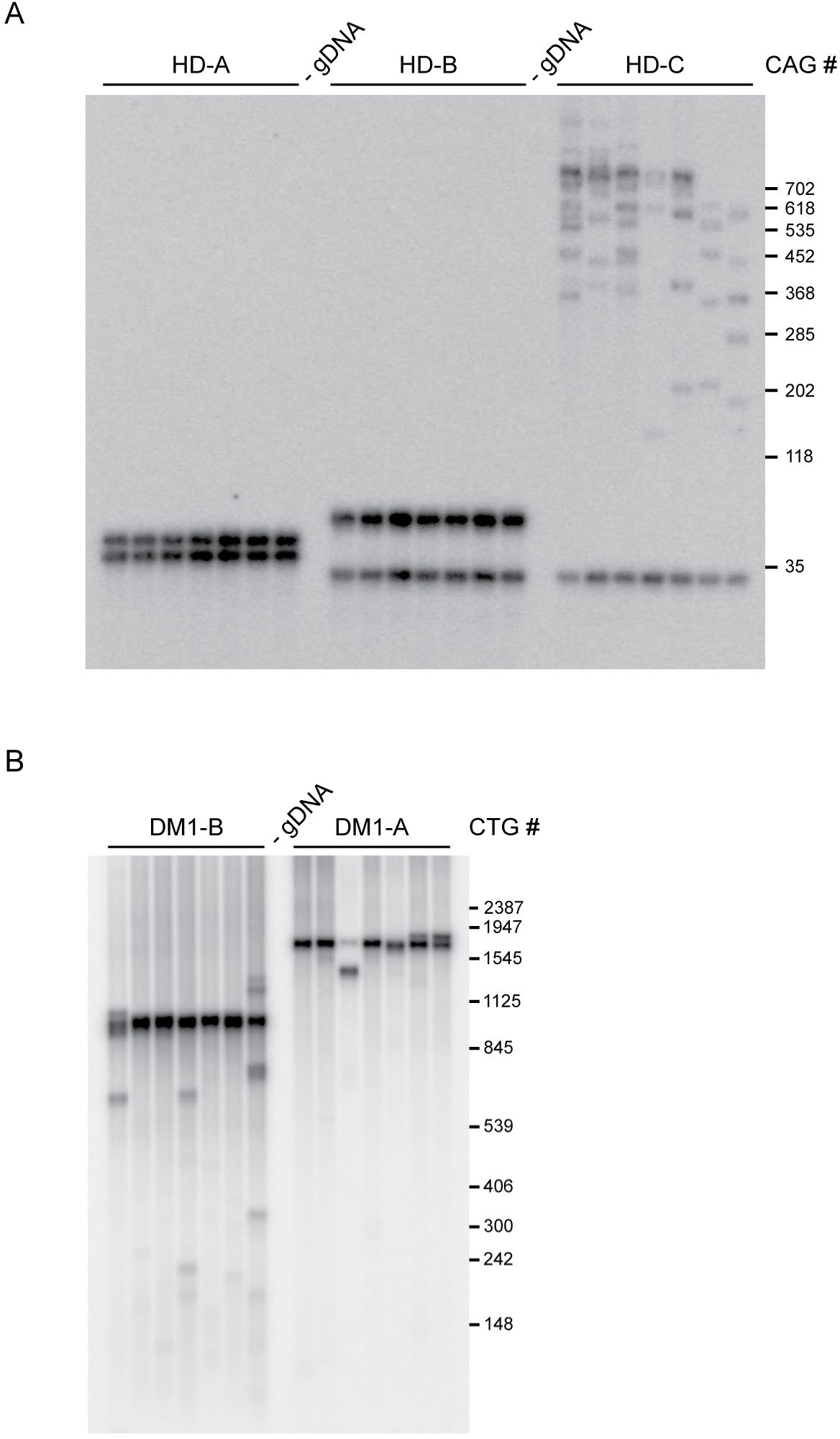
Repeat sizes of HD and DM1 patient cell lines. **(A)** Small-pool PCR of DNA isolated from HD-A, HD-B, and HD-C cells (shown: 1 ng gDNA per PCR). Lanes labeled “-gDNA” are control PCRs with no gDNA. **(B)** Small-pool PCR of DNA isolated from DM1-A and DM1-B cells (shown: 1 ng gDNA per PCR). Lanes labeled “-gDNA” are control PCRs with no gDNA.

**Fig. S2.**
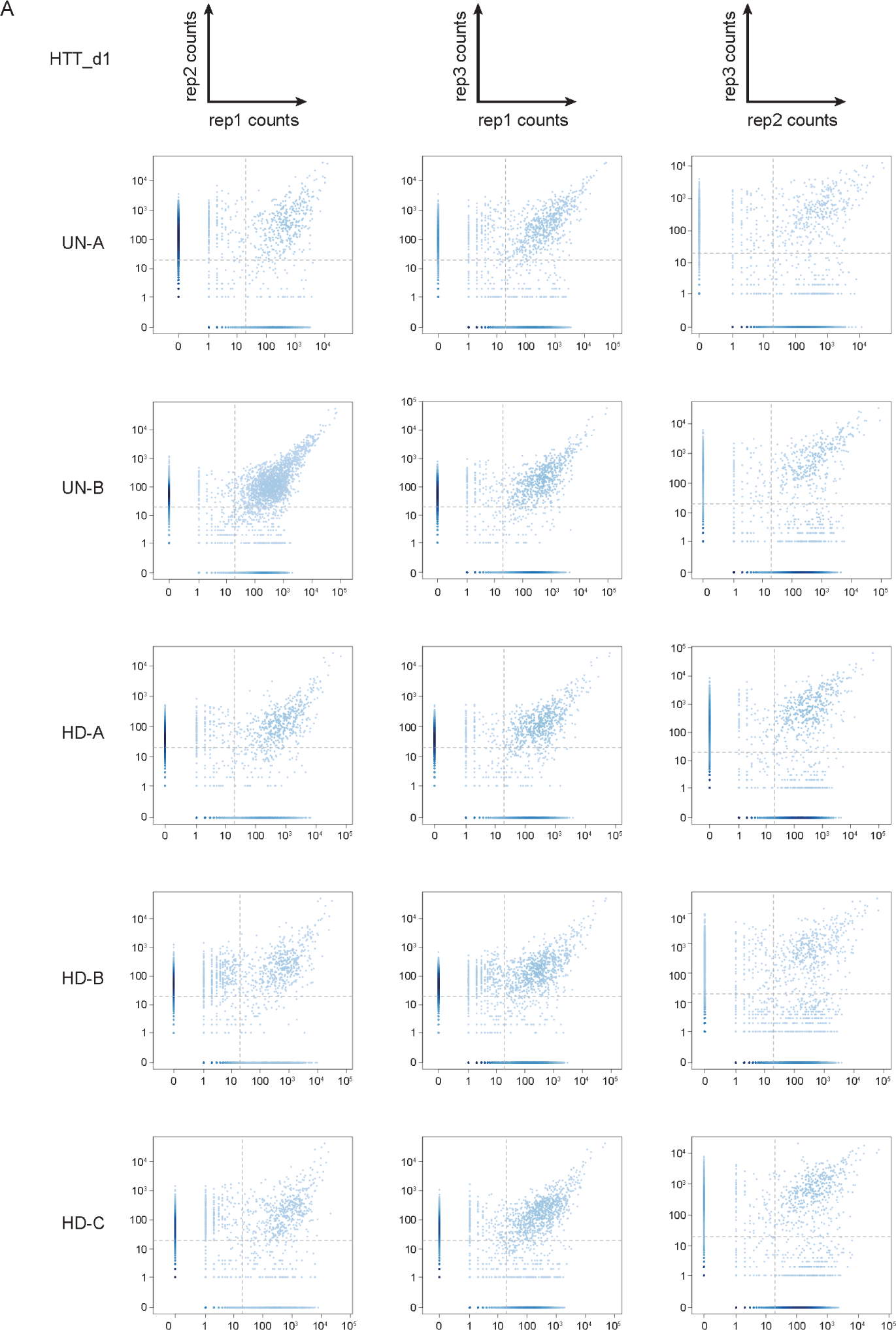
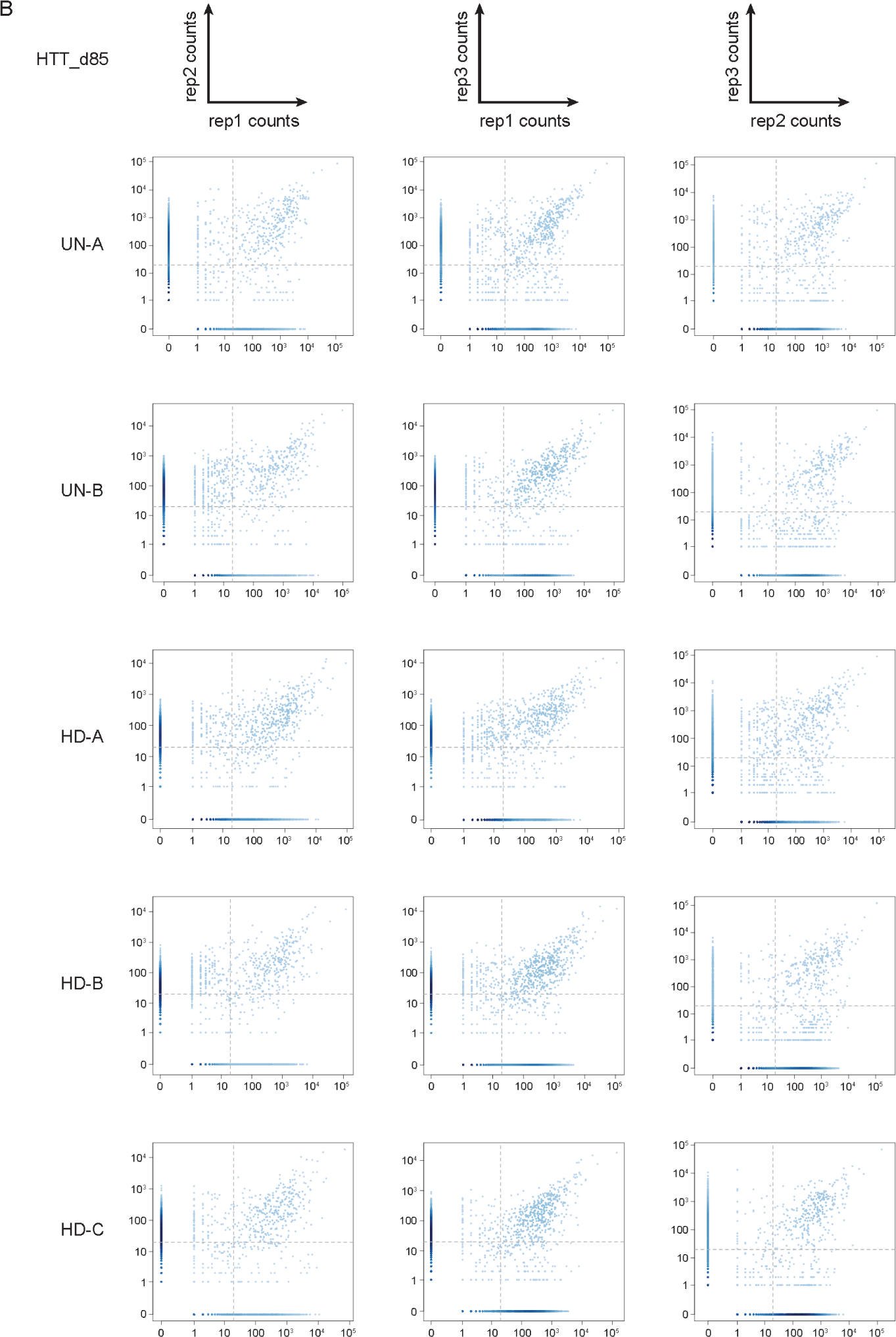
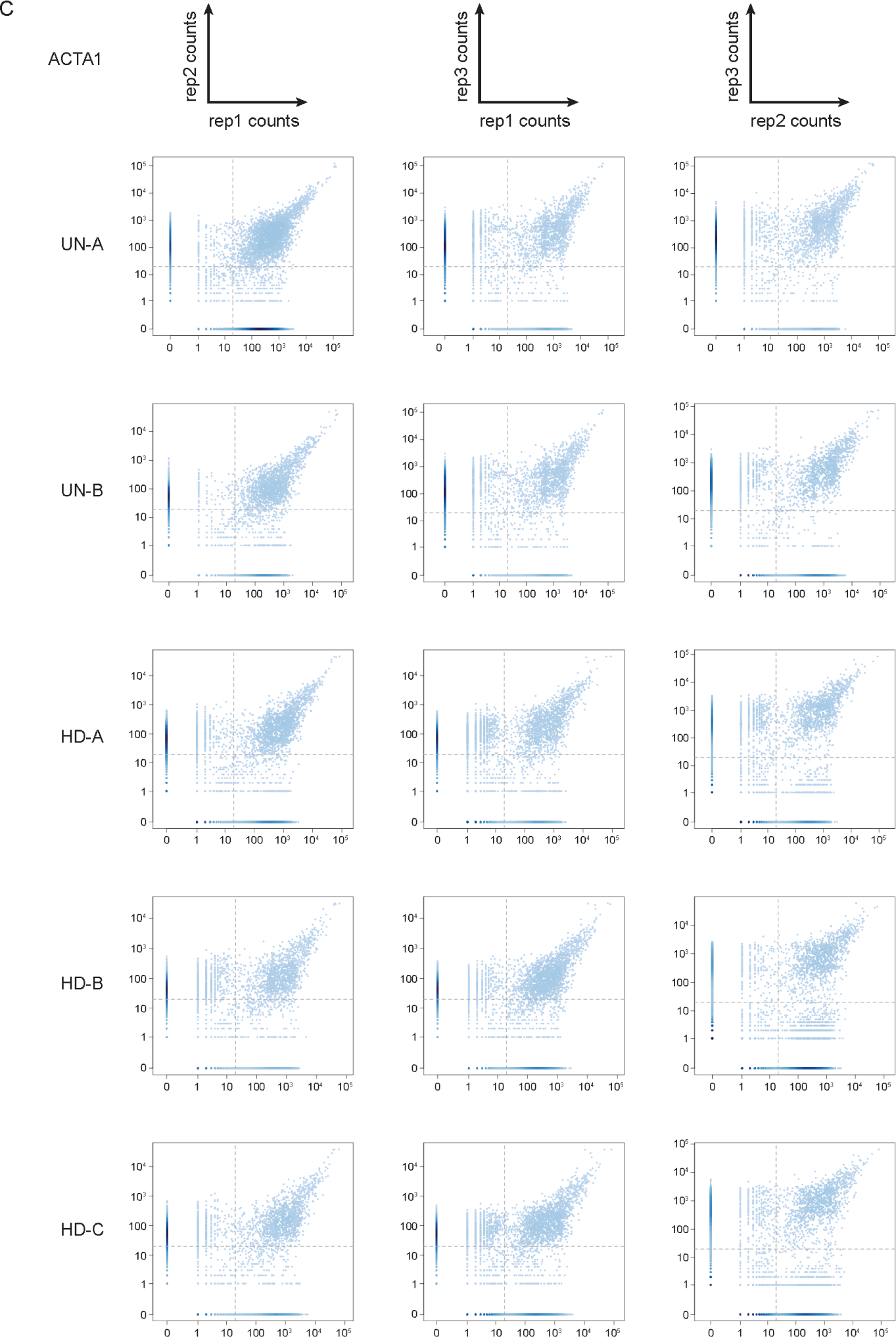
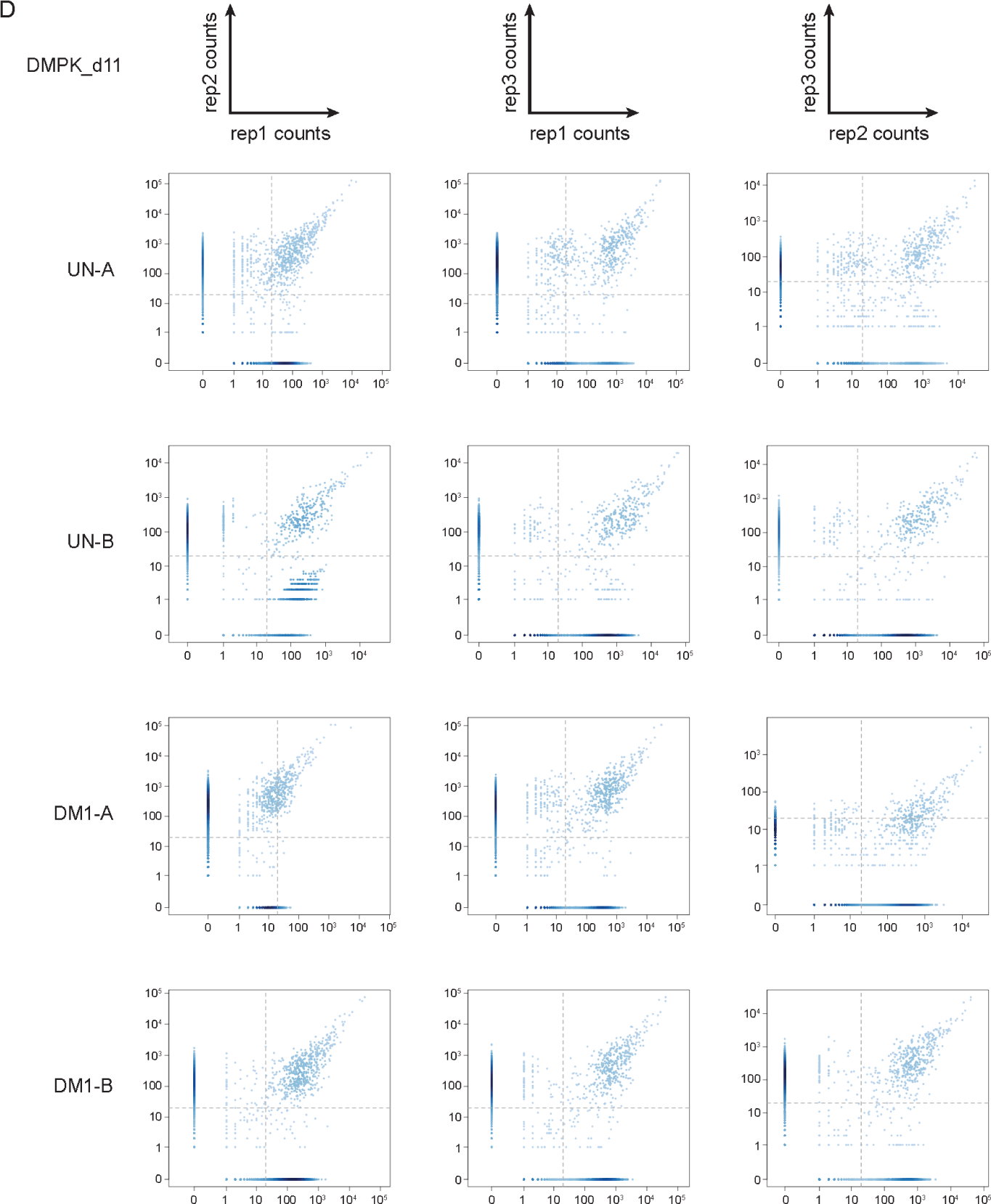
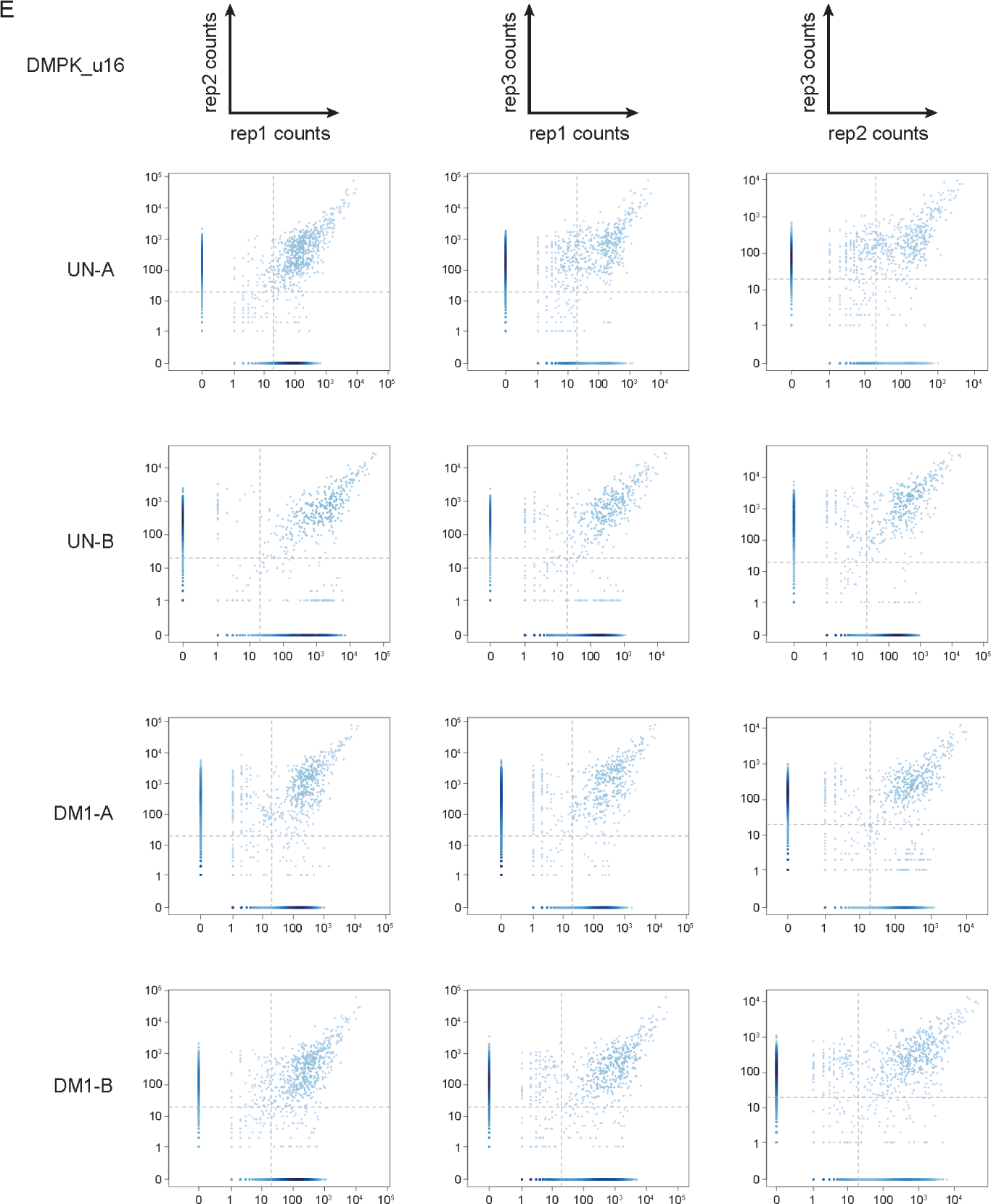
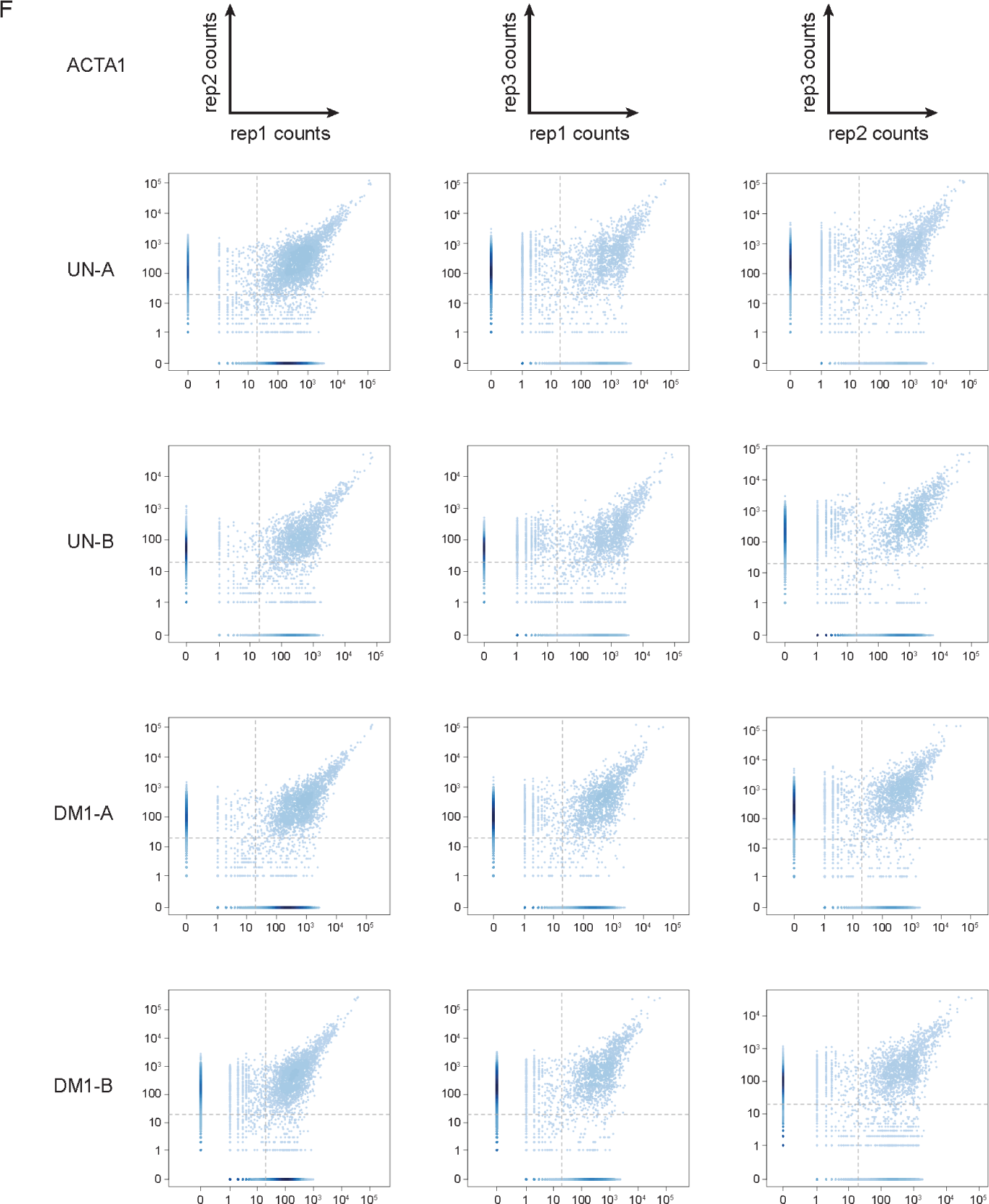
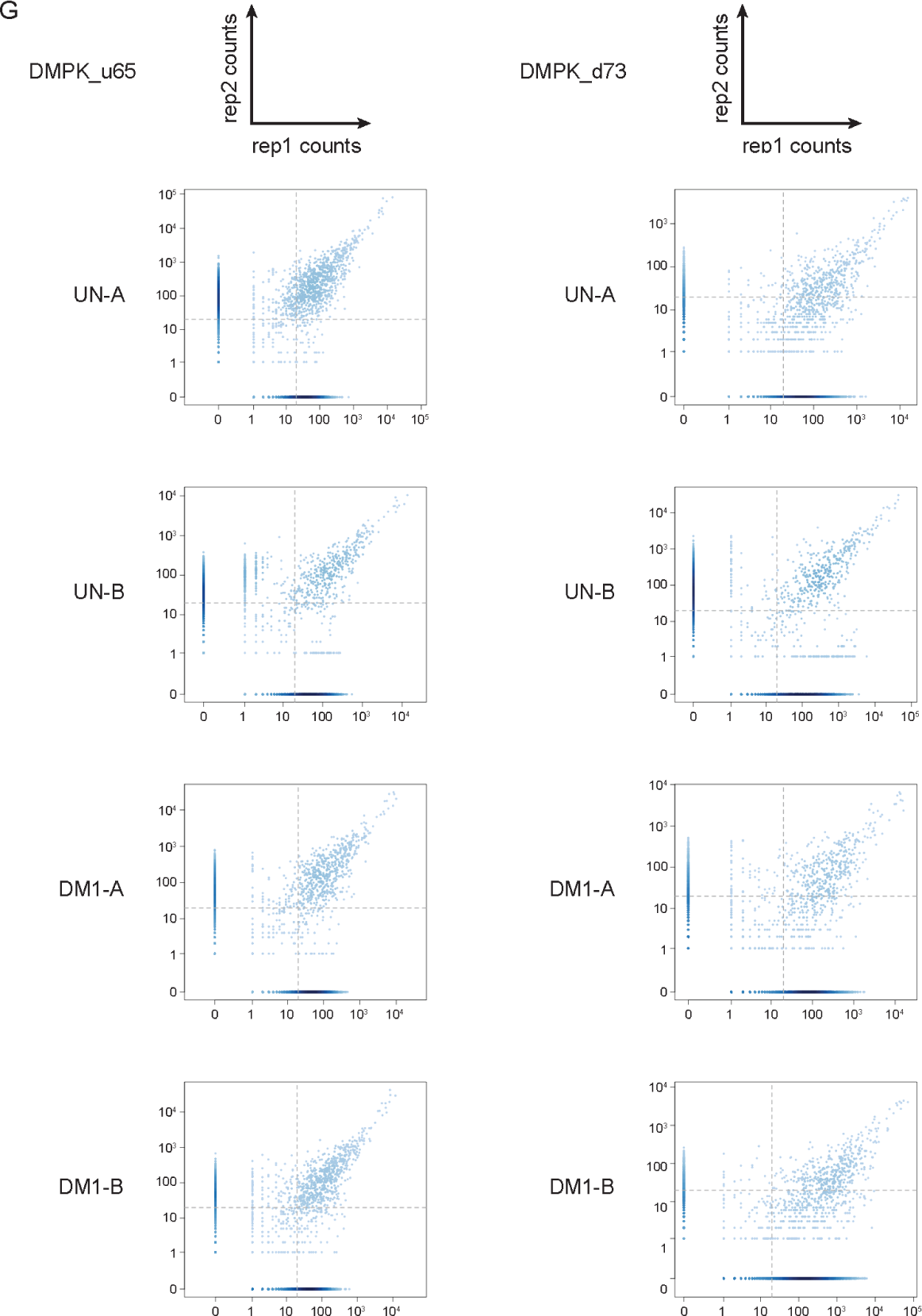
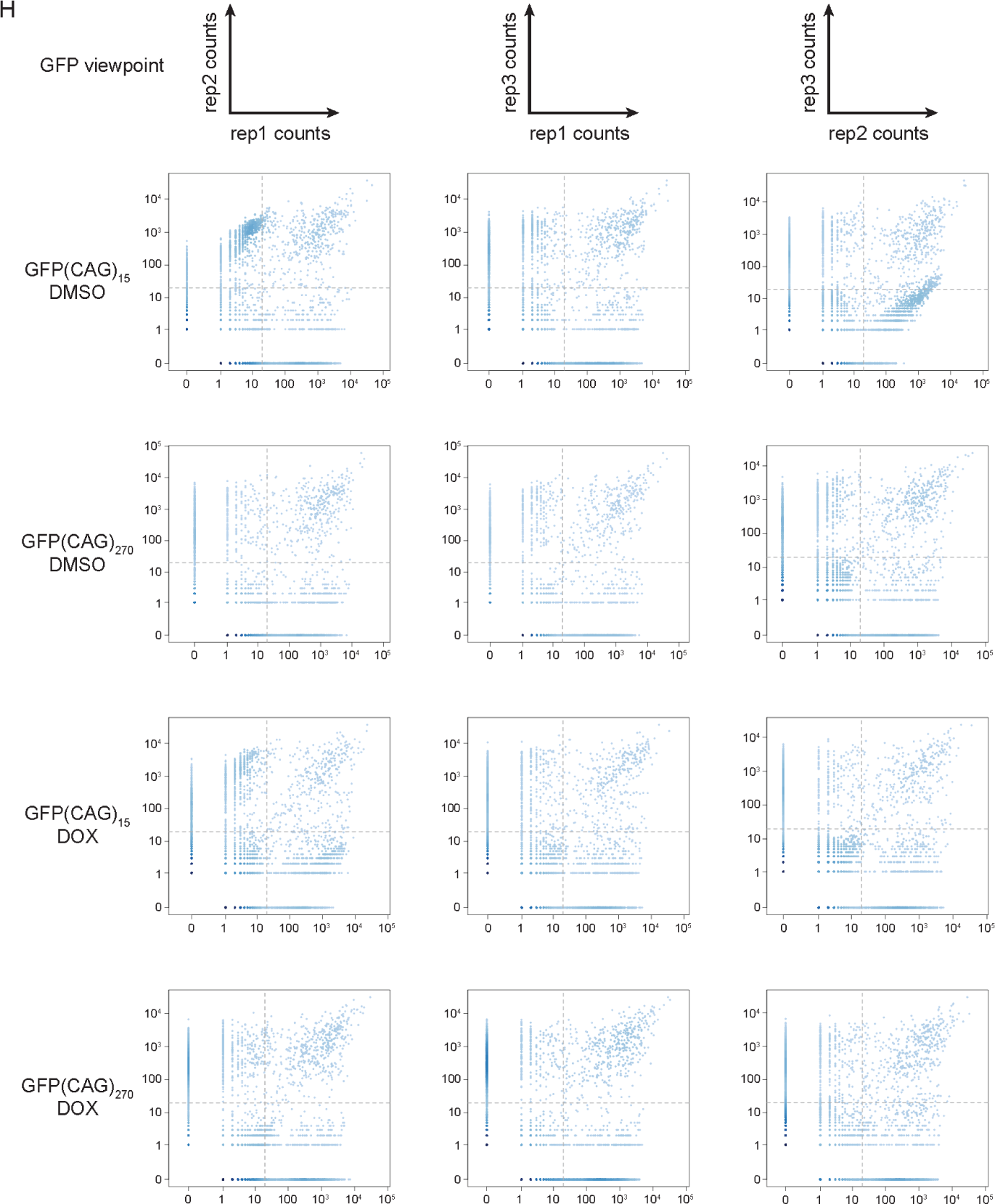
Correlation between 4C library replicates of the *HTT*, *DMPK*, *ACTA1*, and GFP viewpoints. Scatter plots comparing the mapped read counts per 4C fragment between replicates from UN-A, UN-B, HD-A, HD-B, and HD-C cells (A-C); from UN-A, UN-B, DM1-A, and DM1-B cells (D-G); and from GFP(CAG)15 and GFP(CAG)270 (H) separated by viewpoint: **(A)** HTT_d1 viewpoint (1 kb downstream of the CAG repeat), **(B)** HTT_d85 viewpoint (85 kb downstream of the CAG repeat), **(C)** *ACTA1* viewpoint, **(D)** DMPK_d11 viewpoint (11 kb downstream of the CTG repeat), **(E)** DMPK_u16 viewpoint (16 kb upstream of the CTG repeat), **(F)** *ACTA1* viewpoint, **(G)** DMPK_u65 and DMPK_d73 viewpoints (65 kb upstream and 73 kb downstream of the CTG repeat, respectively), and **(H)** GFP viewpoint in GFP(CAG)15 and GFP(CAG)270 cells with transcription off (DMSO) or on (DOX).

**Fig. S3.**
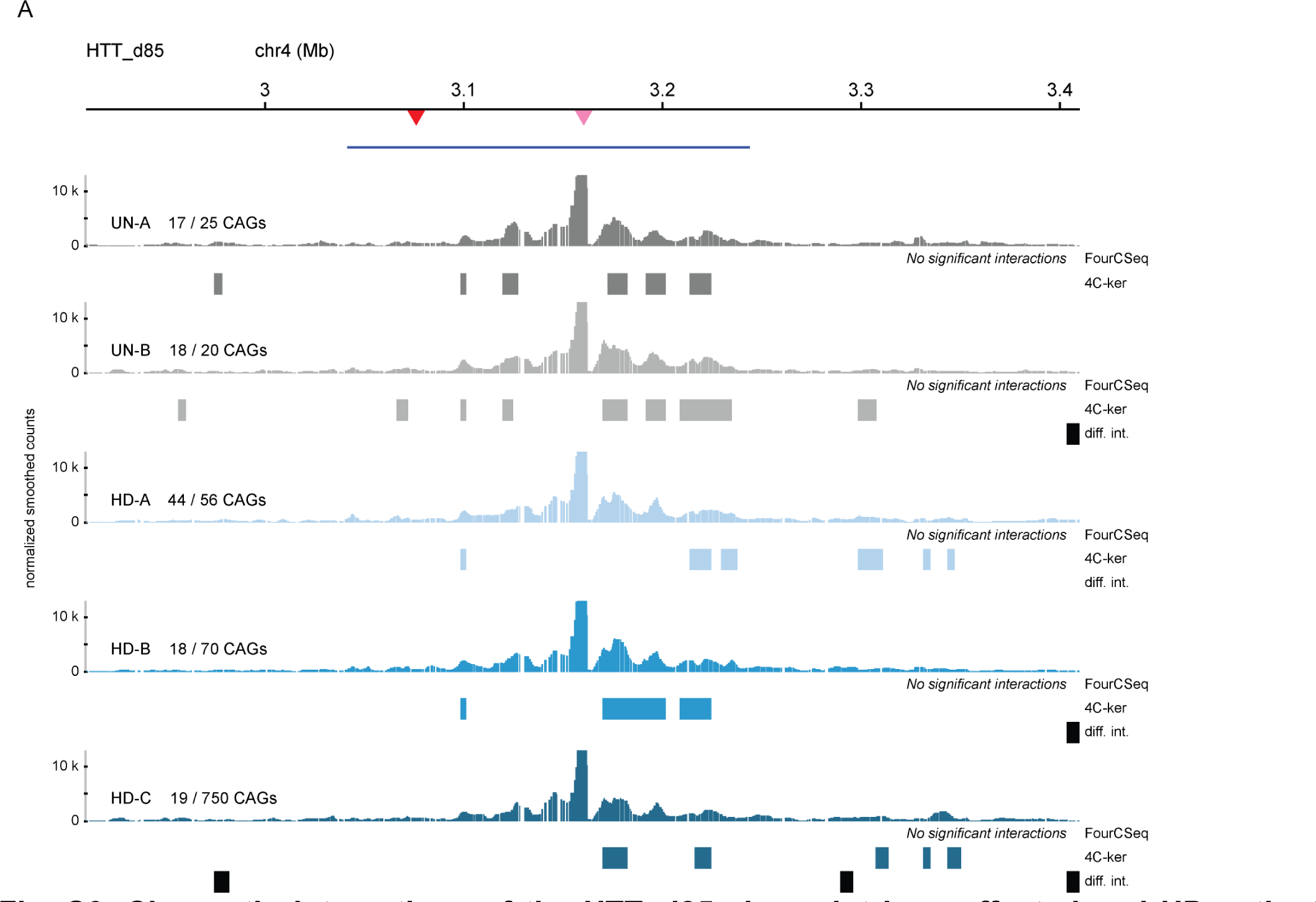
Chromatin interactions of the HTT_d85 viewpoint in unaffected and HD patient cells. **(A)** 4C-seq chromatin interaction profiles from the HTT_d85 viewpoint (85 kb downstream of the CAG repeat) in two unaffected (UN-A and UN-B) and three HD LCLs (HD-A, HD-B, and HD-C). High-interacting regions were called using 4C-ker and significant interactions were called using FourCSeq. Regions of differential interactions are marked with black bars below each 4C-seq track and labeled as “diff. int.”. The top blue bar represents the *HTT* gene. The triangles at the top represent the location of the two *HTT* viewpoints.

**Fig. S4.**
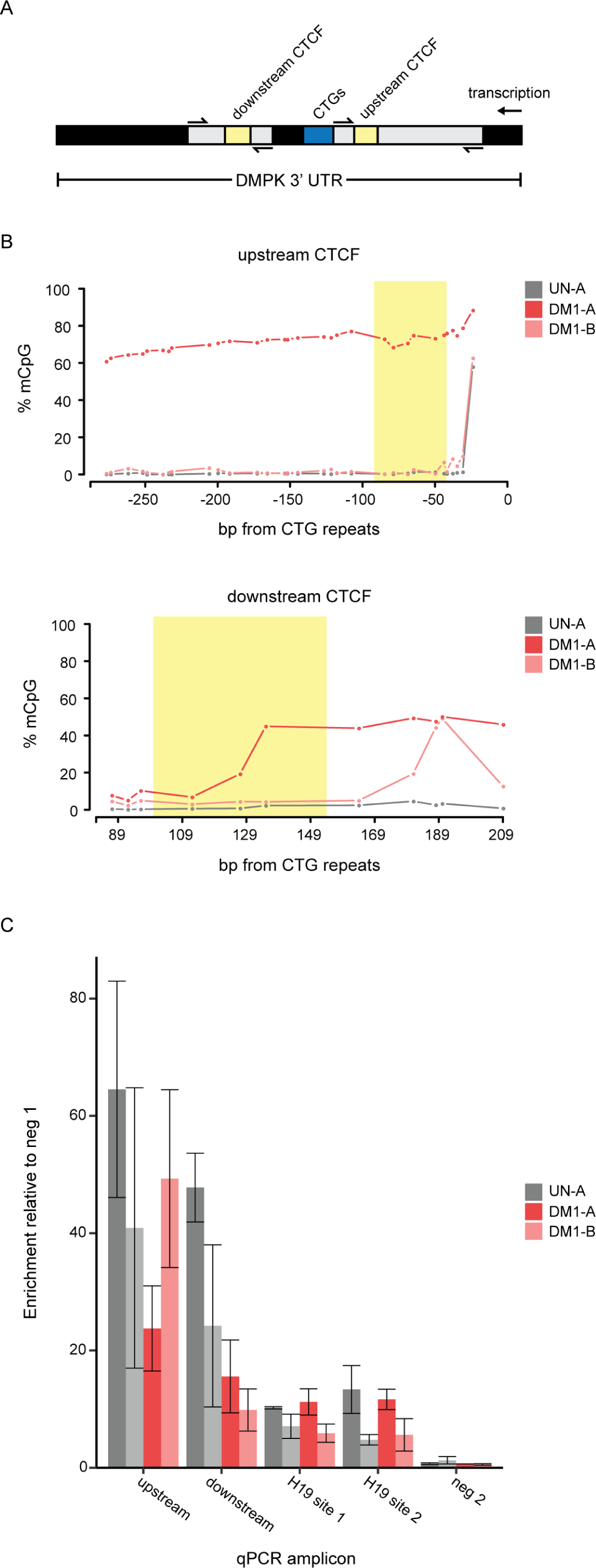
Heterochromatinization of expanded *DMPK* alleles in DM1 patient cells. **(A)** Schematic of the 3’ UTR of *DMPK*. The PCR amplicons used for bisulfite sequencing and CTCF ChIP-qPCR are indicated in grey. The CTCF binding sites are represented by the yellow boxes. **(B)** CpG methylation levels of two CTCF binding sites directly flanking the CTG repeats of *DMPK* using bisulfite-sequencing (top: upstream site; bottom: downstream site). The specific CTCF binding sites are demarcated with yellow bars. **(C)** Chromatin immunoprecipitation of CTCF followed by qPCR at the cognate binding sites found in A and B. This was done in the UN-A, UN-B, DM1-A, and DM1-B LCLs (N=3 for all). CTCF was expected to be bound to the H19 gene, but not at two intergenic sites (neg 1 and neg 2). We normalized the enrichment at each site to neg 1. We found no significant differences when we normalized to the Neg2 site. The error bars represent the standard error of the mean.

**Fig. S5.**
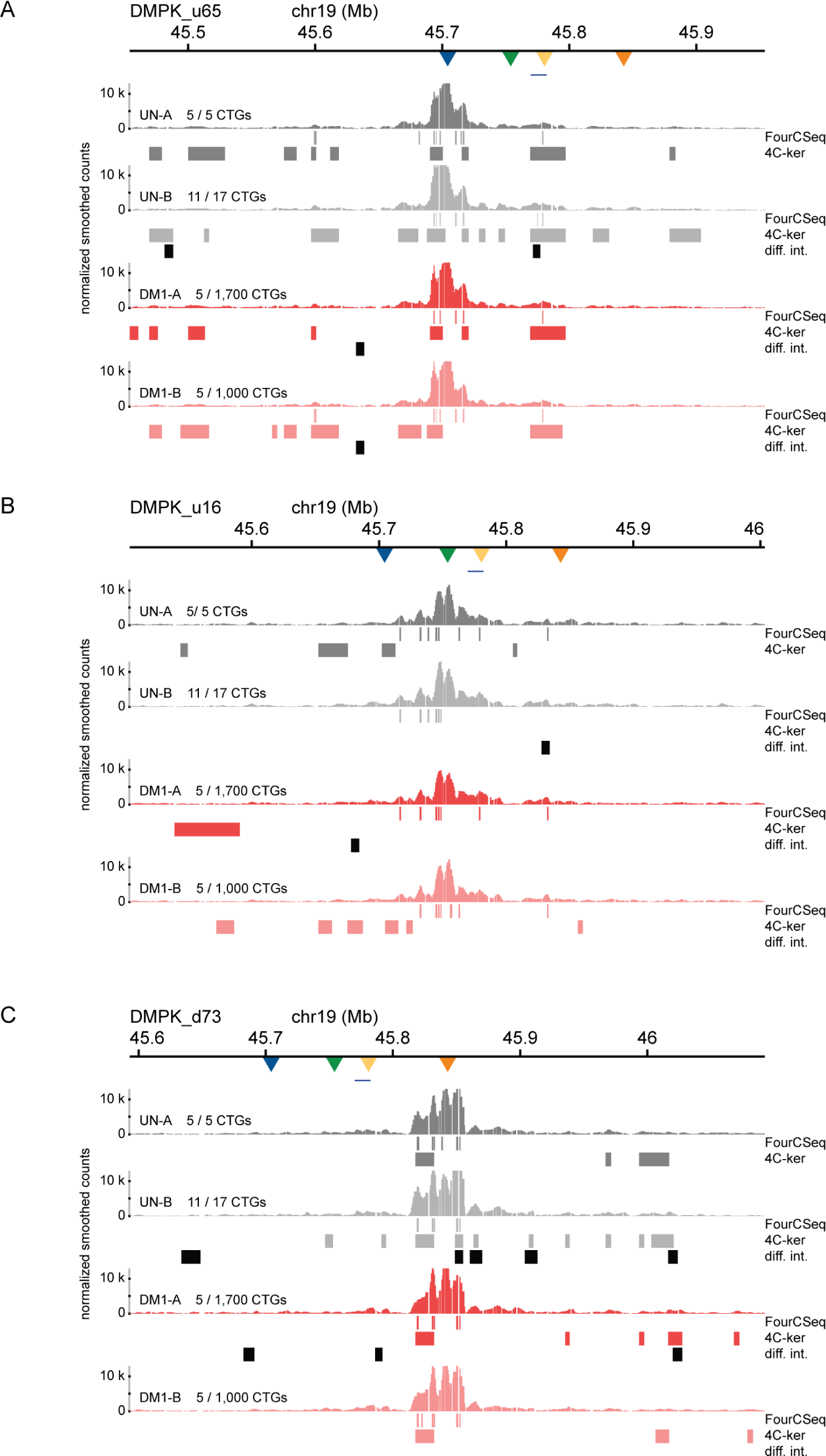
Chromatin interactions at DMPK_u65, DMPK_u16, and DMPK_d73 viewpoints in unaffected and DM1 patient cells. **(A)** 4C-seq chromatin interaction profiles from the DMPK_u65 viewpoint (65 kb upstream of the CAG repeat) in two unaffected (UN-A and UN-B) and two DM1 LCLs (DM1-A and DM1-B). High-interacting regions were called using 4C-ker and significant interactions were called using FourCSeq. Regions of differential interactions are marked with black bars below each 4C-seq track and labeled as “diff. int.”. The top blue bar represents the *DMPK* gene. The triangles at the top represent the location of the four DMPK viewpoints. **(B)** 4C-seq chromatin interaction profiles from the DMPK_u16 viewpoint (16 kb upstream of the CAG repeat) in two unaffected (UN-A and UN-B) and two DM1 LCLs (DM1-A and DM1-B). **(C)** 4C-seq chromatin interaction profiles from the DMPK_d73 viewpoint (73 kb downstream viewpoint in two unaffected (UN-A and UN-B) and two DM1 LCLs (DM1-A and DM1-B).

**Table S1.** Characteristics of cell lines used in this study.

| Cell line | Code | Disease status <sup>1</sup> | Sex <sup>1</sup> | Age at sampling (years) <sup>1</sup> | Repeat locus | # of repeats in wt allele <sup>2</sup> | Sizing method | # of repeats in expanded allele <sup>2</sup> | Sizing method |
| --- | --- | --- | --- | --- | --- | --- | --- | --- | --- |
| GM04604 | UN-A | unaffected | Male | NA | *DMPK* | 5 | Sanger sequencing | 5 | Sanger sequencing |
|  |  |  |  |  | *HTT* | 17 | Sanger sequencing | 25 | Sanger sequencing |
| GM02180 | UN-B | unaffected | Female | 51 | *DMPK* | 11 | Sanger sequencing | 17 | Sanger sequencing |
|  |  |  |  |  | *HTT* | 18 | Sanger sequencing | 20 | Sanger sequencing |
| GM06077 | DM1-A | DM1 | Female | 4 | *DMPK* | 5 | Sanger sequencing | 1700 | sp-PCR |
| GM04648 | DM1-B | DM1 | Male | 23 | *DMPK* | 5 | Sanger sequencing | 1000 | sp-PCR |
| GM02164 | HD-A | HD | Male | 58 | *HTT* | 44 | sp-PCR | 56 | sp-PCR |
| GM03620 | HD-B | HD | Female | 29 | *HTT* | 18 | Sanger sequencing | 70 | sp-PCR |
| GM14044 | HD-C | HD | Male | 16 | *HTT* | 19 | Sanger sequencing | 750 * | sp-PCR |

<sup>1</sup> Information obtained from the Coriell Institute Human Genetic Cell Repository.
<sup>2</sup> For sp-PCR, the number of repeats is an estimation of the modal number of repeats.
NA: not available
sp-PCR: small pool-PCR

**Table S3.** 4C-seq library statistics.

| Sample | # of reads | # of mapped reads | % of total | # of reads mapped to cis chr | % of cis chr reads from total | # of reads mapped to valid 4C frags in cis chr | % of reads in 4C frags from total cis chr reads |
| --- | --- | --- | --- | --- | --- | --- | --- |
| DMPK_u65_GM04604_rep1 | 11'192'157 | 8'116'207 | 72.5 | 2'296'664 | 28.3 | 1'805'386 | 78.6 |
| DMPK_u65_GM04604_rep2 | 1'688'853 | 1'035'697 | 61.3 | 606'020 | 58.5 | 513'147 | 84.7 |
| DMPK_u65_GM02180_rep1 | 1'183'618 | 725'020 | 61.3 | 402'692 | 55.5 | 342'602 | 85.1 |
| DMPK_u65_GM02180_rep2 | 1'461'840 | 888'805 | 60.8 | 466'817 | 52.5 | 406'381 | 87.1 |
| DMPK_u65_GM06077_rep1 | 8'498'158 | 5'535'188 | 65.1 | 1'053'071 | 19 | 908'403 | 86.3 |
| DMPK_u65_GM06077_rep2 | 1'586'296 | 966'005 | 60.9 | 574'031 | 59.4 | 515'578 | 89.8 |
| DMPK_u65_GM04648_rep1 | 8'646'819 | 5'368'775 | 62.1 | 1'045'112 | 19.5 | 942'967 | 90.2 |
| DMPK_u65_GM04648_rep2 | 2'167'574 | 1'221'577 | 56.4 | 668'154 | 54.7 | 588'914 | 88.1 |
| DMPK_u16_GM04604_rep1 | 16'969'035 | 13'541'809 | 79.8 | 2'368'499 | 17.5 | 1'502'637 | 63.4 |
| DMPK_u16_GM04604_rep2 | 8'471'505 | 1'910'469 | 22.6 | 611'665 | 32 | 391'201 | 64 |
| DMPK_u16_GM04604_rep3 | 5'023'436 | 4'174'649 | 83.1 | 325'026 | 7.8 | 214'817 | 66.1 |
| DMPK_u16_GM02180_rep1 | 4'967'776 | 3'272'865 | 65.9 | 1'156'773 | 35.3 | 807'405 | 69.8 |
| DMPK_u16_GM02180_rep2 | 9'791'581 | 6'074'072 | 62 | 2'791'710 | 46 | 1'918'545 | 68.7 |
| DMPK_u16_GM02180_rep3 | 3'499'397 | 2'404'710 | 68.7 | 743'880 | 30.9 | 494'040 | 66.4 |
| DMPK_u16_GM06077_rep1 | 44'300'878 | 31'633'701 | 71.4 | 4'230'357 | 13.4 | 3'136'434 | 74.1 |
| DMPK_u16_GM06077_rep2 | 7'516'556 | 2'615'231 | 34.8 | 791'991 | 30.3 | 512'797 | 64.7 |
| DMPK_u16_GM06077_rep3 | 3'344'272 | 2'426'791 | 72.6 | 693'947 | 28.6 | 436'064 | 62.8 |
| DMPK_u16_GM04648_rep1 | 14'372'890 | 11'367'448 | 79.1 | 1'616'380 | 14.2 | 1'080'070 | 66.8 |
| DMPK_u16_GM04648_rep2 | 5'456'784 | 1'941'577 | 35.6 | 874'656 | 45 | 560'803 | 64.1 |
| DMPK_u16_GM04648_rep3 | 8'354'943 | 5'260'739 | 63 | 2'365'753 | 45 | 1'549'651 | 65.5 |
| DMPK_d11_GM04604_rep1 | 23'059'825 | 10'787'846 | 46.8 | 2'388'136 | 22.1 | 1'782'296 | 74.6 |
| DMPK_d11_GM04604_rep2 | 8'042'866 | 717'825 | 8.9 | 355'974 | 49.6 | 251'617 | 70.7 |
| DMPK_d11_GM04604_rep3 | 6'716'905 | 4'244'006 | 63.2 | 1'416'961 | 33.4 | 988'977 | 69.8 |
| DMPK_d11_GM02180_rep1 | 4'296'953 | 866'253 | 20.2 | 442'187 | 51 | 330'779 | 74.8 |
| DMPK_d11_GM02180_rep2 | 3'308'800 | 894'717 | 27 | 412'986 | 46.2 | 308'579 | 74.7 |
| DMPK_d11_GM02180_rep3 | 7'282'955 | 4'517'085 | 62 | 2'175'116 | 48.2 | 1'518'060 | 69.8 |
| DMPK_d11_GM06077_rep1 | 15'395'452 | 7'880'443 | 51.2 | 2'460'619 | 31.2 | 1'843'809 | 74.9 |
| DMPK_d11_GM06077_rep2 | 774'586 | 109'054 | 14.1 | 56'680 | 52 | 41'488 | 73.2 |
| DMPK_d11_GM06077_rep3 | 6'524'825 | 2'605'272 | 39.9 | 1'097'272 | 42.1 | 773'163 | 70.5 |
| DMPK_d11_GM04648_rep1 | 8'560'029 | 4'972'802 | 58.1 | 1'413'824 | 28.4 | 979'700 | 69.3 |
| DMPK_d11_GM04648_rep2 | 12'454'988 | 1'910'083 | 15.3 | 928'582 | 48.6 | 678'608 | 73.1 |
| DMPK_d11_GM04648_rep3 | 8'259'430 | 3'846'132 | 46.6 | 1'529'463 | 39.8 | 1'063'240 | 69.5 |
| DMPK_d73_GM04604_rep1 | 5'976'375 | 5'457'650 | 91.3 | 325'342 | 6 | 171'443 | 52.7 |
| DMPK_d73_GM04604_rep2 | 4'710'733 | 2'209'035 | 46.9 | 816'530 | 37 | 507'446 | 62.1 |
| DMPK_d73_GM02180_rep1 | 4'278'924 | 2'351'635 | 55 | 781'457 | 33.2 | 512'019 | 65.5 |
| DMPK_d73_GM02180_rep2 | 6'211'002 | 4'265'102 | 68.7 | 1'463'307 | 34.3 | 918'581 | 62.8 |
| DMPK_d73_GM06077_rep1 | 25'744'531 | 24'078'279 | 93.5 | 2'287'837 | 9.5 | 334'267 | 14.6 |
| DMPK_d73_GM06077_rep2 | 5'923'808 | 2'670'770 | 45.1 | 871'927 | 32.6 | 565'589 | 64.9 |
| DMPK_d73_GM04648_rep1 | 8'317'618 | 7'753'683 | 93.2 | 686'800 | 8.9 | 210'742 | 30.7 |
| DMPK_d73_GM04648_rep2 | 13'833'430 | 8'816'046 | 63.7 | 3'351'872 | 38 | 2'184'265 | 65.2 |
| ACTA1_d2_GM04604_rep1 | 17'462'688 | 9'831'359 | 56.3 | 5'908'533 | 60.1 | 4'186'550 | 70.9 |
| ACTA1_d2_GM04604_rep2 | 20'016'202 | 13'201'152 | 66 | 9'429'955 | 71.4 | 6'749'626 | 71.6 |
| ACTA1_d2_GM04604_rep3 | 14'370'212 | 9'150'078 | 63.7 | 6'493'921 | 71 | 4'854'805 | 74.8 |
| ACTA1_d2_GM02180_rep1 | 4'445'588 | 2'894'160 | 65.1 | 2'047'454 | 70.7 | 1'448'565 | 70.7 |
| ACTA1_d2_GM02180_rep2 | 9'724'309 | 6'455'364 | 66.4 | 4'490'505 | 69.6 | 3'238'325 | 72.1 |
| ACTA1_d2_GM02180_rep3 | 14'471'204 | 9'623'984 | 66.5 | 6'832'538 | 71 | 5'111'636 | 74.8 |
| ACTA1_d2_GM06077_rep1 | 11'677'329 | 7'881'675 | 67.5 | 4'502'685 | 57.1 | 3'148'013 | 69.9 |
| ACTA1_d2_GM06077_rep2 | 20'390'273 | 13'386'906 | 65.7 | 9'414'304 | 70.3 | 6'751'808 | 71.7 |
| ACTA1_d2_GM06077_rep3 | 6'528'425 | 4'468'502 | 68.4 | 3'101'465 | 69.4 | 2'186'147 | 70.5 |
| ACTA1_d2_GM04648_rep1 | 28'697'528 | 18'150'768 | 63.2 | 9'891'236 | 54.5 | 7'190'802 | 72.7 |
| ACTA1_d2_GM04648_rep2 | 6'247'015 | 4'135'124 | 66.2 | 2'850'483 | 68.9 | 2'025'028 | 71 |
| ACTA1_d2_GM04648_rep3 | 7'615'653 | 5'226'254 | 68.6 | 3'625'835 | 69.4 | 2'493'472 | 68.8 |
| ACTA1_d2_GM02164_rep1 | 4'982'938 | 3'226'684 | 64.8 | 2'262'970 | 70.1 | 1'598'786 | 70.6 |
| ACTA1_d2_GM02164_rep2 | 14'175'182 | 9'482'669 | 66.9 | 6'432'203 | 67.8 | 4'689'799 | 72.9 |
| ACTA1_d2_GM02164_rep3 | 8'985'756 | 6'149'600 | 68.4 | 4'314'765 | 70.2 | 3'032'024 | 70.3 |
| ACTA1_d2_GM03620_rep1 | 3'392'689 | 2'252'717 | 66.4 | 1'594'521 | 70.8 | 1'125'432 | 70.6 |
| ACTA1_d2_GM03620_rep2 | 10'153'899 | 6'911'566 | 68.1 | 4'682'012 | 67.7 | 3'412'228 | 72.9 |
| ACTA1_d2_GM03620_rep3 | 10'314'868 | 6'898'449 | 66.9 | 4'788'393 | 69.4 | 3'452'374 | 72.1 |
| ACTA1_d2_GM14044_rep1 | 3'864'340 | 2'522'916 | 65.3 | 1'719'045 | 68.1 | 1'218'110 | 70.9 |
| ACTA1_d2_GM14044_rep2 | 13'566'864 | 8'987'115 | 66.2 | 6'029'854 | 67.1 | 4'498'420 | 74.6 |
| ACTA1_d2_GM14044_rep3 | 10'730'254 | 7'441'130 | 69.3 | 5'128'011 | 68.9 | 3'821'275 | 74.5 |
| HTT_d1_GM04604_rep1 | 4'212'566 | 2'457'650 | 58.3 | 1'404'970 | 57.2 | 1'103'125 | 78.5 |
| HTT_d1_GM04604_rep2 | 10'749'708 | 1'832'850 | 17.1 | 888'698 | 48.5 | 639'666 | 72 |
| HTT_d1_GM04604_rep3 | 5'921'707 | 3'530'799 | 59.6 | 2'010'544 | 56.9 | 1'556'448 | 77.4 |
| HTT_d1_GM02180_rep1 | 2'641'610 | 1'678'631 | 63.5 | 975'311 | 58.1 | 687'885 | 70.5 |
| HTT_d1_GM02180_rep2 | 4'261'764 | 2'775'669 | 65.1 | 1'331'970 | 48 | 972'583 | 73 |
| HTT_d1_GM02180_rep3 | 6'713'195 | 3'878'698 | 57.8 | 2'112'225 | 54.5 | 1'644'256 | 77.8 |
| HTT_d1_GM02164_rep1 | 1'547'564 | 1'006'519 | 65 | 550'667 | 54.7 | 378'050 | 68.7 |
| HTT_d1_GM02164_rep2 | 6'426'843 | 4'114'843 | 64 | 2'049'276 | 49.8 | 1'475'404 | 72 |
| HTT_d1_GM02164_rep3 | 4'561'062 | 2'649'147 | 58.1 | 1'529'524 | 57.7 | 1'141'625 | 74.6 |
| HTT_d1_GM03620_rep1 | 2'799'272 | 1'795'212 | 64.1 | 1'031'035 | 57.4 | 704'440 | 68.3 |
| HTT_d1_GM03620_rep2 | 5'475'190 | 3'487'162 | 63.7 | 1'558'114 | 44.7 | 1'137'993 | 73 |
| HTT_d1_GM03620_rep3 | 7'098'155 | 3'663'907 | 51.6 | 2'006'314 | 54.8 | 1'576'201 | 78.6 |
| HTT_d1_GM14044_rep1 | 2'595'315 | 1'668'384 | 64.3 | 945'665 | 56.7 | 659'221 | 69.7 |
| HTT_d1_GM14044_rep2 | 6'212'554 | 3'953'292 | 63.6 | 1'666'102 | 42.1 | 1'189'773 | 71.4 |
| HTT_d1_GM14044_rep3 | 5'199'603 | 2'755'357 | 53 | 1'459'570 | 53 | 1'128'283 | 77.3 |
| HTT_d85_GM04604_rep1 | 3'626'944 | 2'088'253 | 57.6 | 1'300'447 | 62.3 | 850'403 | 65.4 |
| HTT_d85_GM04604_rep2 | 4'387'441 | 2'691'863 | 61.4 | 1'062'983 | 39.5 | 637'080 | 59.9 |
| HTT_d85_GM04604_rep3 | 4'022'177 | 2'243'116 | 55.8 | 1'343'423 | 59.9 | 882'695 | 65.7 |
| HTT_d85_GM02180_rep1 | 2'884'811 | 1'333'135 | 46.2 | 773'017 | 58 | 467'647 | 60.5 |
| HTT_d85_GM02180_rep2 | 4'805'661 | 3'283'731 | 68.3 | 1'132'457 | 34.5 | 719'942 | 63.6 |
| HTT_d85_GM02180_rep3 | 3'937'446 | 2'278'255 | 57.9 | 1'335'882 | 58.6 | 909'086 | 68.1 |
| HTT_d85_GM02164_rep1 | 2'065'583 | 773'630 | 37.5 | 440'336 | 56.9 | 279'660 | 63.5 |
| HTT_d85_GM02164_rep2 | 4'271'456 | 2'674'960 | 62.6 | 1'320'008 | 49.3 | 846'436 | 64.1 |
| HTT_d85_GM02164_rep3 | 3'365'643 | 1'762'199 | 52.4 | 1'048'467 | 59.5 | 674'809 | 64.4 |
| HTT_d85_GM03620_rep1 | 3'134'531 | 654'436 | 20.9 | 397'137 | 60.7 | 245'017 | 61.7 |
| HTT_d85_GM03620_rep2 | 3'232'273 | 2'024'049 | 62.6 | 978'716 | 48.4 | 577'818 | 59 |
| HTT_d85_GM03620_rep3 | 4'074'032 | 2'290'232 | 56.2 | 1'356'058 | 59.2 | 898'455 | 66.3 |
| HTT_d85_GM14044_rep1 | 2'057'402 | 798'621 | 38.8 | 453'565 | 56.8 | 276'126 | 60.9 |
| HTT_d85_GM14044_rep2 | 5'576'017 | 2'543'284 | 45.6 | 956'326 | 37.6 | 583'814 | 61 |
| HTT_d85_GM14044_rep3 | 3'538'845 | 2'077'028 | 58.7 | 1'191'091 | 57.3 | 772'206 | 64.8 |
| GFP_CAG_15_DMSO_rep1 | 7'048'343 | 2'833'417 | 40.2 | 1'700'523 | 60 | 1'278'717 | 75.2 |
| GFP_CAG_15_DMSO_rep2 | 7'487'213 | 2'887'212 | 38.6 | 1'638'799 | 56.8 | 1'284'684 | 78.4 |
| GFP_CAG_15_DMSO_rep3 | 6'936'377 | 2'769'339 | 39.9 | 1'464'325 | 52.9 | 1'172'916 | 80.1 |
| GFP_CAG_15_DOX_rep1 | 11'304'727 | 4'562'245 | 40.4 | 2'182'322 | 47.8 | 1'705'839 | 78.2 |
| GFP_CAG_15_DOX_rep2 | 8'452'926 | 3'124'374 | 37 | 1'540'855 | 49.3 | 1'198'057 | 77.8 |
| GFP_CAG_15_DOX_rep3 | 7'631'549 | 2'833'020 | 37.1 | 1'466'250 | 51.8 | 1'157'955 | 79 |
| GFP_CAG_270_DMSO_rep1 | 8'320'728 | 3'099'916 | 37.3 | 1'907'976 | 61.5 | 1'512'580 | 79.3 |
| GFP_CAG_270_DMSO_rep2 | 6'840'505 | 2'694'631 | 39.4 | 1'575'886 | 58.5 | 1'229'060 | 78 |
| GFP_CAG_270_DMSO_rep3 | 7'338'988 | 2'673'561 | 36.4 | 1'569'149 | 58.7 | 1'225'397 | 78.1 |
| GFP_CAG_270_DOX_rep1 | 9'301'866 | 3'679'030 | 39.6 | 2'078'639 | 56.5 | 1'620'884 | 78 |
| GFP_CAG_270_DOX_rep2 | 8'353'878 | 3'289'609 | 39.4 | 1'735'082 | 52.7 | 1'333'717 | 76.9 |
| GFP_CAG_270_DOX_rep3 | 7'577'290 | 3'045'799 | 40.2 | 1'588'724 | 52.2 | 1'267'573 | 79.8 |

**Table S4.**
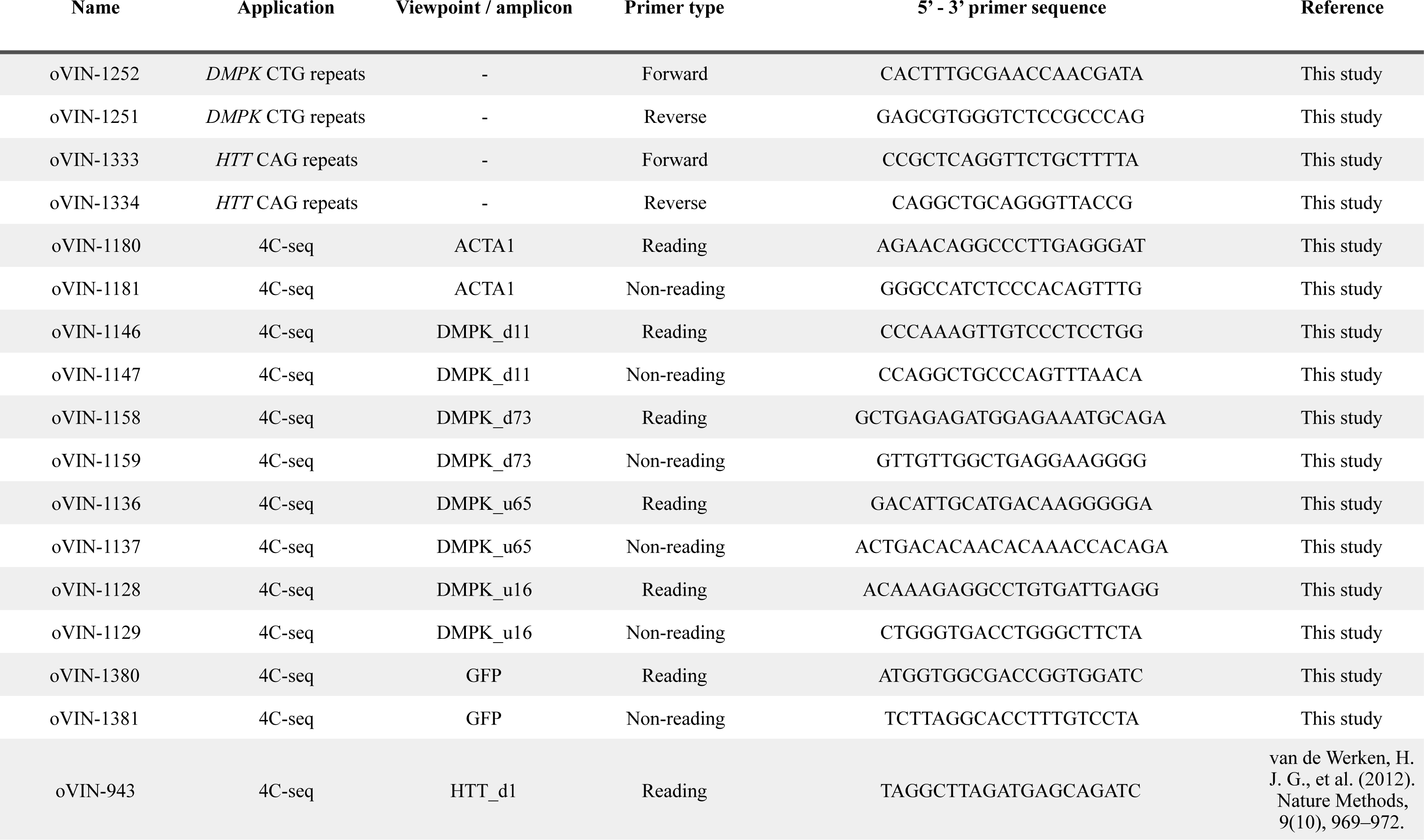
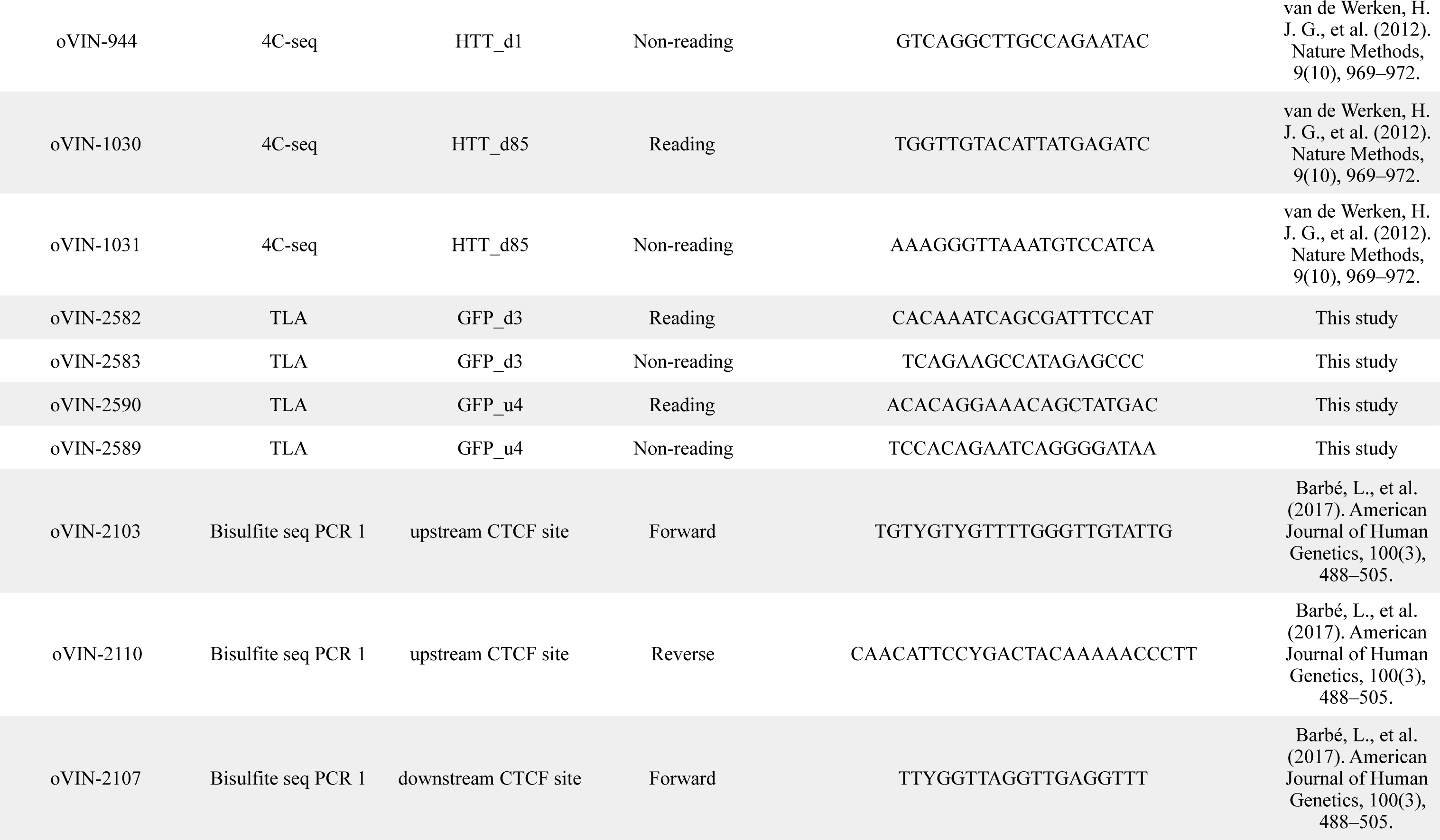
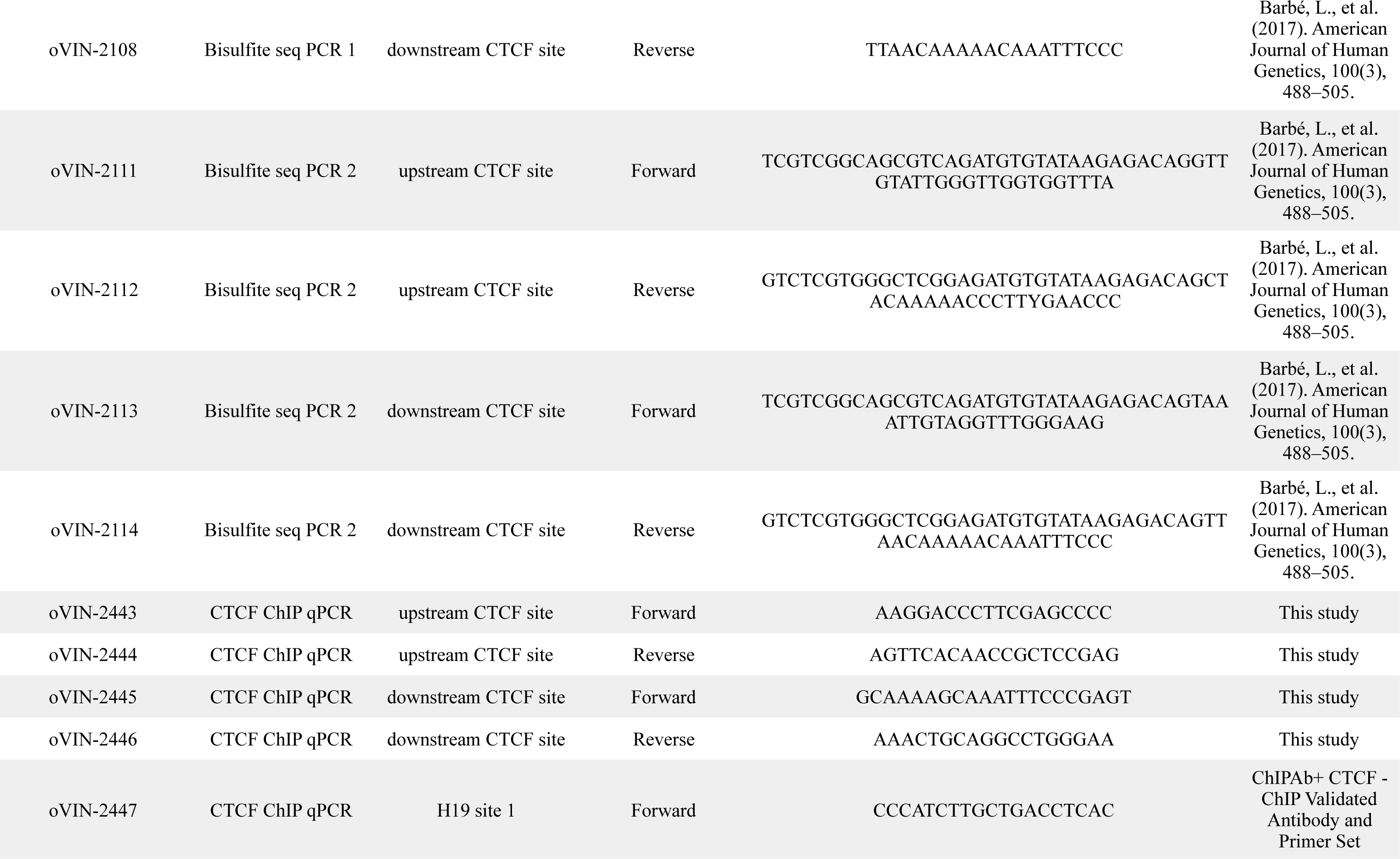
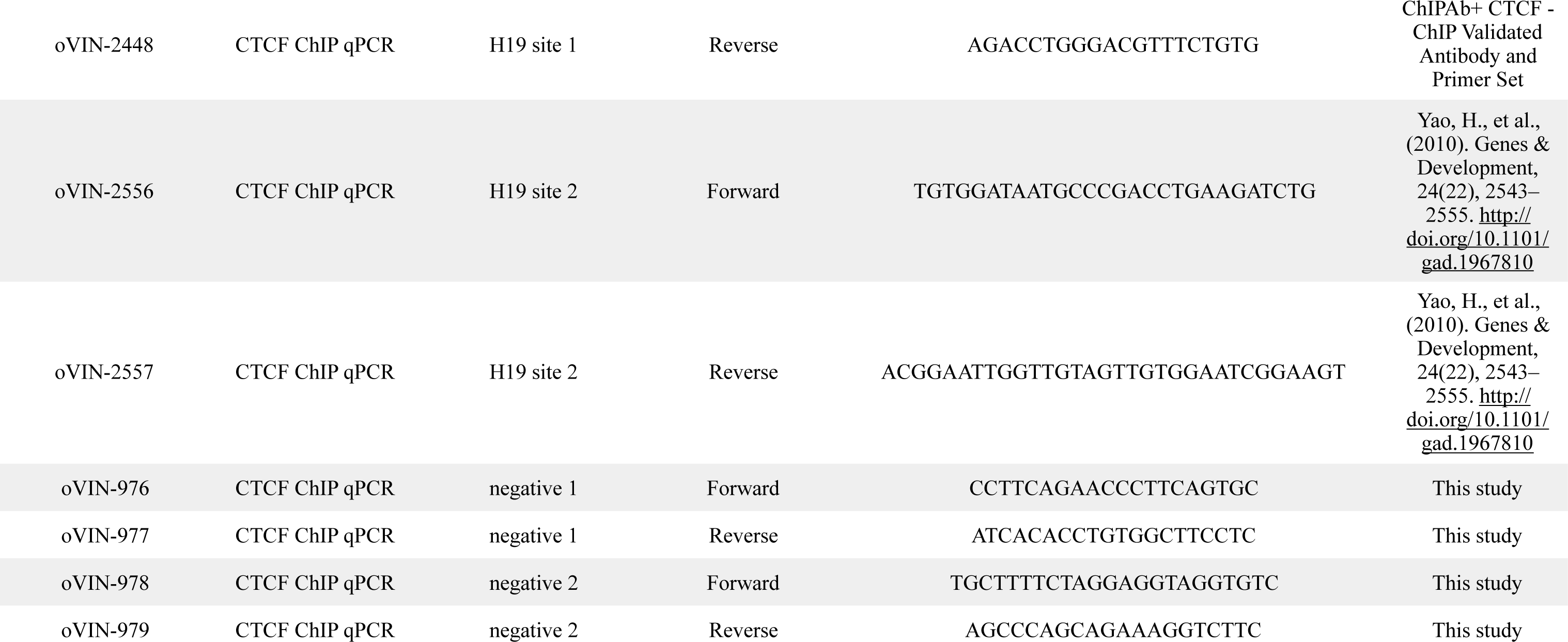
PCR primers used in this study.

## References

1. J. H. Gibcus, J. Dekker, The hierarchy of the 3D genome. Mol. Cell. 49, 773–782 (2013).

2. W. A. Bickmore, B. van Steensel, Genome architecture: domain organization of interphase chromosomes. Cell. 152, 1270–1284 (2013).

3. A. Pombo, N. Dillon, Three-dimensional genome architecture: players and mechanisms. Nat. Rev. Mol. Cell Biol. 16, 245–257 (2015).

4. D. G. Lupiáñez, M. Spielmann, S. Mundlos, Breaking TADs: How Alterations of Chromatin Domains Result in Disease. Trends Genet. 32, 225–237 (2016).

5. M. Spielmann, D. G. Lupiáñez, S. Mundlos, Structural variation in the 3D genome. Nat. Rev. Genet. 19, 453–467 (2018).

6. V. Dion, J. H. Wilson, Instability and chromatin structure of expanded trinucleotide repeats. Trends Genet. 25, 288–297 (2009).

7. A. López Castel, J. D. Cleary, C. E. Pearson, Repeat instability as the basis for human diseases and as a potential target for therapy. Nat. Rev. Mol. Cell Biol. 11, 165–170 (2010).

8. H. T. Orr, H. Y. Zoghbi, Trinucleotide repeat disorders. Annu. Rev. Neurosci. 30, 575–621 (2007).

9. C. Sandi, M. Sandi, S. A. Virmouni, S. Al-Mahdawi, M. A. Pook, Epigenetic-based therapies for Friedreich ataxia. Front Genet. 5 (2014), doi:10.3389/fgene.2014.00165.

10. D. Colak et al., Promoter-Bound Trinucleotide Repeat mRNA Drives Epigenetic Silencing in Fragile X Syndrome. Science. 343, 1002–1005 (2014).

11. E. Tabolacci, P. Chiurazzi, Epigenetics, fragile X syndrome and transcriptional therapy. Am. J. Med. Genet. A. 161A, 2797–2808 (2013).

12. P. K. Chan et al., Heterochromatinization induced by GAA-repeat hyperexpansion in Friedreich’s ataxia can be reduced upon HDAC inhibition by vitamin B3. Hum. Mol. Genet. 22, 2662–2675 (2013).

13. A. López Castel et al., Expanded CTG repeat demarcates a boundary for abnormal CpG methylation in myotonic dystrophy patient tissues. Hum. Mol. Genet. 20, 1–15 (2011).

14. D. Kumari, K. Usdin, The distribution of repressive histone modifications on silenced FMR1 alleles provides clues to the mechanism of gene silencing in fragile X syndrome. Hum. Mol. Genet. 19, 4634–4642 (2010).

15. D. Kumari, K. Usdin, Chromatin remodeling in the noncoding repeat expansion diseases. J. Biol. Chem. 284, 7413–7417 (2009).

16. . D. H. Cho et al., Antisense transcription and heterochromatin at the DM1 CTG repeats are constrained by CTCF. Mol. Cell. 20, 483–489 (2005).

17. G. N. Filippova et al., CTCF-binding sites flank CTG/CAG repeats and form a methylation-sensitive insulator at the DM1 locus. Nat. Genet. 28, 335–343 (2001).

18. A. J. Verkerk et al., Identification of a gene (FMR-1) containing a CGG repeat coincident with a breakpoint cluster region exhibiting length variation in fragile X syndrome. Cell. 65, 905–914 (1991).

19. Y. H. Fu, D. Kuhl, A. Pizzuti, M. Pieretti, J. S. Sutcliffe, Variation of the CGG repeat at the fragile X site results in genetic instability: resolution of the Sherman paradox. Cell. 67, 1047–1058 (1991).

20. I. Oberlé et al., Instability of a 550-base pair DNA segment and abnormal methylation in fragile X syndrome. Science. 252, 1097–1102 (1991).

21. J. P. Fryns, A. Kleczkowska, E. Kubien, H. Vandenberghe, Cytogenetic Findings in Moderate and Severe Mental-Retardation - a Study of an Institutionalized Population of 1991 Patients. Acta Paediatr Scand Suppl. 313, 1–& (1984).

22. V. Campuzano et al., Friedreich’s ataxia: autosomal recessive disease caused by an intronic GAA triplet repeat expansion. Science. 271, 1423–1427 (1996).

23. N. Gheldof, T. M. Tabuchi, J. Dekker, The active FMR1 promoter is associated with a large domain of altered chromatin conformation with embedded local histone modifications. Proc. Natl. Acad. Sci. U.S.A. 103, 12463–12468 (2006).

24. R. Pietrobono et al., Molecular dissection of the events leading to inactivation of the FMR1 gene. Hum. Mol. Genet. 14, 267–277 (2005).

25. B. Coffee, F. P. Zhang, S. Ceman, S. T. Warren, D. Reines, Histone modifications depict an aberrantly heterochromatinized FMR1 gene in fragile X syndrome. Am. J. Hum. Genet. 71, 923–932 (2002).

26. B. Coffee, F. Zhang, S. T. Warren, D. Reines, Acetylated histones are associated with FMR1 in normal but not fragile X-syndrome cells. Nat. Genet. 22, 98–101 (1999).

27. I. De Biase, Y. K. Chutake, P. M. Rindler, S. I. Bidichandani, Epigenetic silencing in Friedreich ataxia is associated with depletion of CTCF (CCCTC-binding factor) and antisense transcription. PLoS ONE. 4, e7914 (2009).

28. S. Al-Mahdawi et al., The Friedreich ataxia GAA repeat expansion mutation induces comparable epigenetic changes in human and transgenic mouse brain and heart tissues. Hum. Mol. Genet. 17, 735–746 (2008).

29. E. Soragni et al., Long intronic GAA*TTC repeats induce epigenetic changes and reporter gene silencing in a molecular model of Friedreich ataxia. Nucleic Acids Res. 36, 6056–6065 (2008).

30. E. Greene, L. Mahishi, A. Entezam, D. Kumari, K. Usdin, Repeat-induced epigenetic changes in intron 1 of the frataxin gene and its consequences in Friedreich ataxia. Nucleic Acids Res. 35, 3383–3390 (2007).

31. J. H. Sun et al., Disease-Associated Short Tandem Repeats Co-localize with Chromatin Domain Boundaries. Cell, 1–31 (2018).

32. J. R. Brouwer, A. Huguet, A. Nicole, A. Munnich, G. Gourdon, Transcriptionally Repressive Chromatin Remodelling and CpG Methylation in the Presence of Expanded CTG-Repeats at the DM1 Locus. J Nucleic Acids. 2013, 567435–16 (2013).

33. T. R. Klesert et al., Mice deficient in Six5 develop cataracts: implications for myotonic dystrophy. Nat. Genet. 25, 105–109 (2000).

34. A. D. Otten, S. J. Tapscott, Triplet repeat expansion in myotonic dystrophy alters the adjacent chromatin structure. Proc. Natl. Acad. Sci. U.S.A. 92, 5465–5469 (1995).

35. X. A. Su, V. Dion, S. M. Gasser, C. H. Freudenreich, Regulation of recombination at yeast nuclear pores controls repair and triplet repeat stability. Genes Dev. 29, 1006–1017 (2015).

36. H. J. G. van de Werken et al., Robust 4C-seq data analysis to screen for regulatory DNA interactions. Nat. Methods. 9, 969–972 (2012).

37. M. Simonis et al., Nuclear organization of active and inactive chromatin domains uncovered by chromosome conformation capture-on-chip (4C). Nat. Genet. 38, 1348–1354 (2006).

38. F. A. Klein et al., FourCSeq: analysis of 4C sequencing data. Bioinformatics. 31, 3085–3091 (2015).

39. R. Raviram et al., 4C-ker: A Method to Reproducibly Identify Genome-Wide Interactions Captured by 4C-Seq Experiments. PLoS Comput Biol. 12, e1004780–23 (2016).

40. L. Barbé et al., CpG Methylation, a Parent-of-Origin Effect for Maternal-Biased Transmission of Congenital Myotonic Dystrophy. Am. J. Hum. Genet. 100, 488–505 (2017).

41. C. Cinesi, L. E. N. Aeschbach, Bin Yang, V. Dion, Contracting CAG/CTG repeats using the CRISPR-Cas9 nickase. Nat Commun. 7, 1–10 (2016).

42. B. A. Santillan, C. Moye, D. Mittelman, J. H. Wilson, GFP-based fluorescence assay for CAG repeat instability in cultured human cells. PLoS ONE. 9, e113952 (2014).

43. P. J. P. de Vree et al., Targeted sequencing by proximity ligation for comprehensive variant detection and local haplotyping. Nat. Biotechnol. 32, 1019–1025 (2014).

44. A. R. Barutcu, B. J. Blencowe, J. L. Rinn, Differential contribution of steady-state RNA and active transcription in chromatin organization. EMBO Rep., e48068 (2019).

45. D. Racko, F. Benedetti, J. Dorier, A. Stasiak, Transcription-induced supercoiling as the driving force of chromatin loop extrusion during formation of TADs in interphase chromosomes. Nucleic Acids Res. 46, 1648–1660 (2018).

46. T. B. Le, M. T. Laub, Transcription rate and transcript length drive formation of chromosomal interaction domain boundaries. EMBO J. 35, 1582–1595 (2016).

47. F. Le Dily et al., Distinct structural transitions of chromatin topological domains correlate with coordinated hormone-induced gene regulation. Genes Dev. 28, 2151–2162 (2014).

48. M. Merkenschlager, E. P. Nora, CTCF and Cohesin in Genome Folding and Transcriptional Gene Regulation. Annu Rev Genomics Hum Genet. 17, 17–43 (2016).

49. R. Adihe Lokanga, X.-N. Zhao, A. Entezam, K. Usdin, X inactivation plays a major role in the gender bias in somatic expansion in a mouse model of the fragile X-related disorders: implications for the mechanism of repeat expansion. Hum. Mol. Genet. 23, 4985–4994 (2014).

50. D. Wohrle, U. Salat, H. Hameister, W. Vogel, P. Steinbach, Demethylation, reactivation, and destabilization of human fragile X full-mutation alleles in mouse embryocarcinoma cells. Am. J. Hum. Genet. 69, 504–515 (2001).

51. C. T. McMurray, Mechanisms of trinucleotide repeat instability during human development. Nat. Rev. Genet. 11, 786–799 (2010).

52. G. De Michele et al., Parental gender, age at birth and expansion length influence GAA repeat intergenerational instability in the X25 gene: pedigree studies and analysis of sperm from patients with Friedreich’s ataxia. Hum. Mol. Genet. 7, 1901–1906 (1998).

53. A. R. Barutcu, P. G. Maass, J. P. Lewandowski, C. L. Weiner, J. L. Rinn, A TAD boundary is preserved upon deletion of the CTCF-rich Firre locus. Nat Commun. 9, 1444 (2018).

54. Y. Ghavi-Helm et al., Highly rearranged chromosomes reveal uncoupling between genome topology and gene expression. Nat. Genet. 144, 327 (2019).

55. V. Dion, Y. Lin, L. Hubert, R. A. Waterland, J. H. Wilson, Dnmt1 deficiency promotes CAG repeat expansion in the mouse germline. Hum. Mol. Genet. 17, 1306–1317 (2008).

56. E. Splinter, E. de Wit, H. J. G. van de Werken, P. Klous, W. de Laat, Determining long-range chromatin interactions for selected genomic sites using 4C-seq technology: from fixation to computation. Methods. 58, 221–230 (2012).

57. F. P. A. David et al., HTSstation: a web application and open-access libraries for high-throughput sequencing data analysis. PLoS ONE. 9, e85879 (2014).

58. D. Noordermeer et al., The dynamic architecture of Hox gene clusters. Science. 334, 222–225 (2011).

59. B. Langmead, S. L. Salzberg, Fast gapped-read alignment with Bowtie 2. Nat. Methods. 9, 357–359 (2012).

60. C. Walter, D. Schuetzmann, F. Rosenbauer, M. Dugas, Benchmarking of 4C-seq pipelines based on real and simulated data. Bioinformatics. 46, e91. (2019).

61. A. McKenna et al., The Genome Analysis Toolkit: a MapReduce framework for analyzing next-generation DNA sequencing data. Genome Res. 20, 1297–1303 (2010).

62. H. Li, A statistical framework for SNP calling, mutation discovery, association mapping and population genetical parameter estimation from sequencing data. Bioinformatics. 27, 2987–2993 (2011).

63. H. Li, R. Durbin, Fast and accurate long-read alignment with Burrows-Wheeler transform. Bioinformatics. 26, 589–595 (2010).

